# RNF25 Ubiquitin E3 activity Safeguards Genome Integrity by Modulating DNA Replication, Transcription and Translation

**DOI:** 10.64898/2026.09.21.753208

**Authors:** Emily Soto-Hidalgo, Carmen Espejo-Serrano, Ricardo de Arellano, Lourdes González-Vinceiro, María Marugán-Cardiel, Iria Domínguez-García, Ana R. Hortal, María Luisa Mateos-Martín, Néstor García-Rodríguez, Raimundo Freire, Gonzalo Millán-Zambrano, Román González-Prieto

**Affiliations:** Andalusian Centre for Regenerative Medicine and Molecular Biology (CABIMER), Universidad de Sevilla-CSIC-Universidad Pablo de Olavide-Junta de Andalucía, Sevilla, Spain; Departamento de Biología Celular, Facultad de Biología, Universidad de Sevilla, Sevilla, Spain; Departamento de Genética, Facultad de Biología, Universidad de Sevilla, Sevilla, Spain; Instituto de Tecnologías Biomédicas, Centro de Investigaciones Biomédicas de Canarias, Facultad de Medicina, Campus Ciencias de la Salud, Universidad de La Laguna, 38200, Tenerife, Spain; Universidad Fernando Pessoa Canarias, Santa María de Guía, 35450, Gran Canaria, Spain; Institute of Biomedicine of Seville, IBiS/Hospital Universitario Virgen del Rocío/CSIC/Universidad de Sevilla, Proteomics Facility, Sevilla, Spain

## Abstract

The human genome encodes several hundreds of ubiquitin E3 enzymes, most of which have poorly understood biological roles. In search for novel ubiquitin E3 enzymes involved in DNA damage tolerance, we identified RNF25 as a candidate to have a role in DNA replication stress tolerance. Under stress conditions, RNF25 translocates to the nucleus in a cGAS-dependent manner, and loss of RNF25 ubiquitin E3 activity leads to the accumulation of replication-dependent ssDNA gaps. Combining functional assays with with mass-spectrometry based proteomics approaches such as TULIP2, iPOND-MS and TurboID, we found that, mechanistically, RNF25 promotes the stability of the replication fork at Transcription-Replication Conflicts by the ubiquitin-mediated clearance of RAD18, mono-ubiquitinated PCNA, H2B-K120ub and RECQL, among others. Lack of RNF25 ubiquitin E3 activity promotes the occurance of Transcription-Replication Conflicts and destabilizes reversed replication forks, enabling re-priming and accumulation of ssDNA gaps behind the replication fork, ultimately compromising genome integrity. Overall, RNF25 regulates DNA replication, transcription and translation in an ubiquitin-based goldilocks.

## INTRODUCTION

Ubiquitination, namely, the covalent attachment of ubiquitin, a 76 amino acids long polypeptide, to acceptor lysines in target proteins, is one of the most common post-translational modifications (PTMs) in cells. The ubiquitination reaction is performed by an enzymatic cascade consisting of E1, E2 and E3 enzymes, being the E3 enzymes responsible for substrate specificity. More than 500 putative ubiquitin E3 enzymes are coded in the human genome. Among them, the Really Interesting New Gene (RING) E3 enzymes catalyze the transfer of the ubiquitin moiety from the E2 enzyme directly to the substrate without intermediate binding of ubiquitin to the E3. Thus, for the RING-type ubiquitin E3 enzymes, while the E3 enzyme determines substrate specificity, it is the E2 enzyme what dictates the type of ubiquitin chain linkage. For most of them, E2s, E3s and substrates, the physiological relevance and consequences remain poorly understood (Reviewed in (*1*)).

The emergence of CRISPR/Cas9 technology has enabled the performance of genome wide genetic screens and the identification of genes which deficiency or activation confer synthetic lethality or viability in combination with different treatments or gene mutations (*2–4*). Consistently, these screens have helped identifying new genes involved in different cellular processes, including members of the ubiquitination cascade (*5–8*).

In attempt to find novel ubiquitin E3 enzymes with a role in DNA damage tolerance processes, we searched in genome-wide studies for RING-type E3 which deficiency caused a growth disadvantage after treatment with agents that cause the formation of DNA adducts, and selected RNF25 as a potential relevant candidate in the tolerance pathways to DNA adduct formation. Consistently, a relevant role for RNF25 in the tolerance to ribosomal stress has been recently described (*7*) and, importantly, an E3 activity independent function for RNF25 in DNA replication stability has been proposed (*9*). Although, mechanistically, the involvement of RNF25 in the maintenance of replication fork stability remains poorly understood.

Here, we studied the nucleus-cytoplasmic dynamics of RNF25 in response to adduct-inducing agents and ribosomal stress. Applying mass spectrometry-based proteomics approaches we obtained mechanistical insight of RNF25 ubiquitin E3 activity and its relevance in DNA replication, transcription and translation. Together, our results indicate that a ubiquitin-based mechanism that includes the RNF25 E3 activity to coordinate the DNA replication, transcription and translation processes exist.

## RESULTS

### RNF25 translocate to the nucleus in a cGAS-dependent manner

Aiming to identify novel ubiquitin E3 enzymes involved in DNA replication, we sought a genome-wide CRISPR/Cas9 screen by Daniel Durocheŕs laboratory (*2*) for proteins which present sensitivity to genotoxic agents that promote the formation of DNA adducts which represent a barrier for replication fork progression. Specifically, we focused only in RING (Really Interesting New Gene) Finger proteins (RNFs) (Supplementary Figure 1A). We noticed that RNF25 was among the most sensitive RNFs to Benzopyrene Diol Epoxide (BPDE), UltraViolet (UV) light, Methyl Methane Sulfonate (MMS) or the PARP inhibitor Olaparib, although this sensitivity was relatively mild in a genome wide context. (Supplementary Figure 1B).

RNF25 had been mainly described to be involved in cytoplasmic processes such as translation quality control (*10, 11*), NF-KB signaling, or protein turnover (*12–15*). Thus, as changes in localization could be important in its function, we decided to investigate the subcellular localization of RNF25 in response to different genotoxic reagents by immunofluorescence. However, both a commercial polyclonal against the C-term of RNF25 and an additional one raised against the N-term of RNF25 failed to detect endogenous RNF25 by immunofluorescence (data not shown). Thus, first we generated a CRISPR/Cas9 knockout mutant for RNF25 (RNF25-KO) in P53-deficient RPE1 cells which was tested for complete knockout by sequencing and immunoblotting with two independent anti-RNF25 antibodies (Supplementary Figure 1C-D). Next, we complemented the RNF25-KO cells with a RNF25-GFP construct expressed at close to endogenous levels (Supplementary Figure 1E). Clonogenic assays (Supplementary Figure 2) did not show any significant sensitivity to the different DNA damaging agents for which RNF25-KO cells showed mild sensitivity showed in (*2*).

Nevertheless, we investigated the localization of RNF25 in response to different genotoxic agents such as MMS or UV light (Figure 1A). In unperturbed conditions, RNF25 localized both in the nucleus and the cytoplasm, consistent with its association the ribosomes (*10, 11*). Interestingly, RNF25 translocated to the nucleus in response to DNA damage induced by MMS and UV light. As both MMS and UV can cause damage on the RNA(*16–19*), and RNF25 has been described to participate in the ribotoxic stress response(*10, 11*), we hypothesized that ribosome stalling was promoting RNF25 translocation to the nucleus. Therefore, we treated the cells with the translation inhibitor Cycloheximide (CHX) and analyzed the cells by immunofluorescence (Figure 1B). Accordingly, CHX treatment also promoted RNF25 translocation to the nucleus.

**Figure 1.**
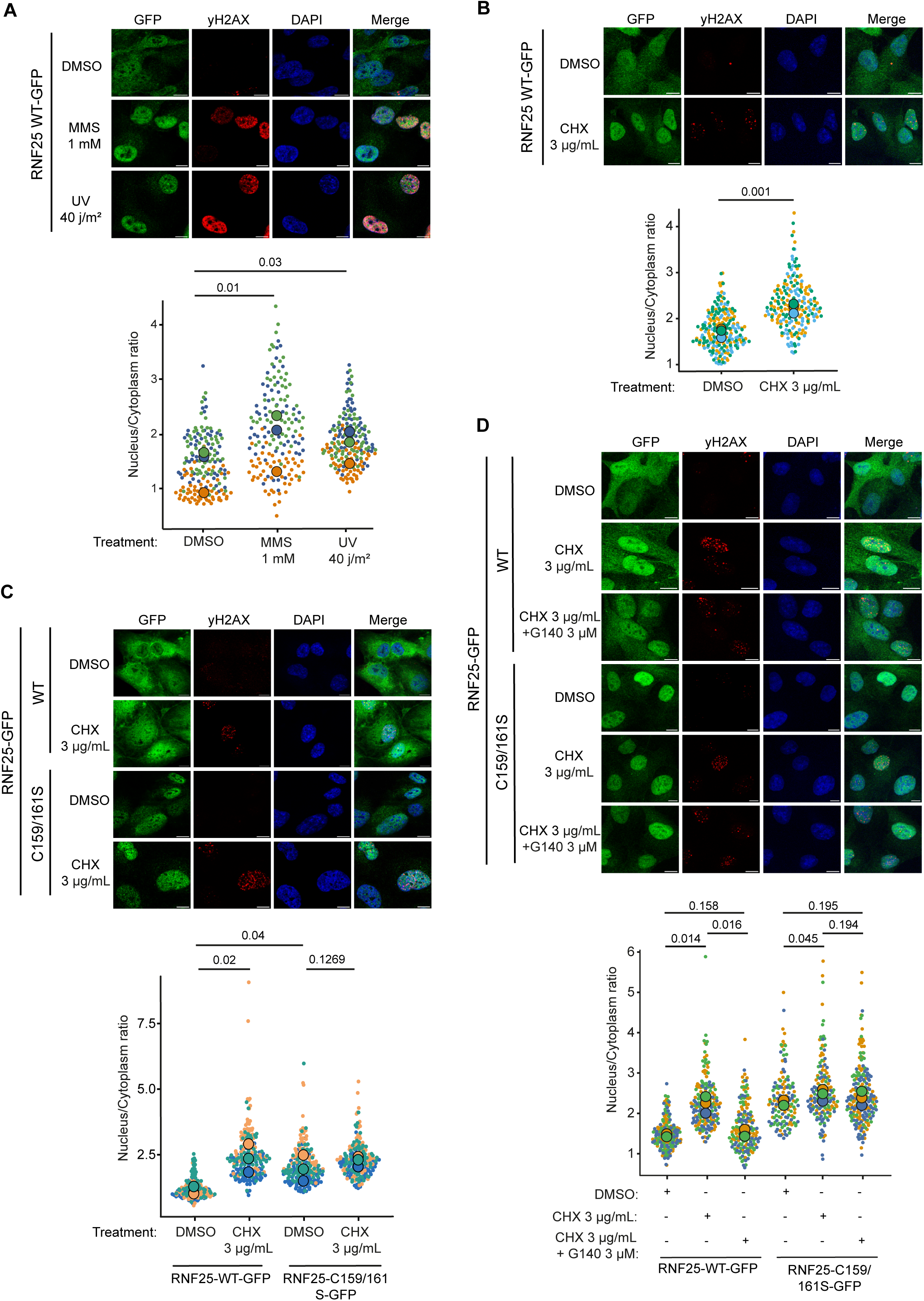
RNF25 translocates to nuclei in response to translational stress. **(A, B)** Immunofluorescence analysis of RNF25 KO cells rescued with RNF25-WT-GFP, treated or not with 1 mM Methyl methanesulfonate (MMS) for 3 hours, irradiated with Ultraviolet (UV) light at 40 j/m^2^ with a 5-hour recovery time, and **(B)** with 3 µg/mL cycloheximide (CHX) for 3 hours. **(C-D)** Immunofluorescence analysis of RNF25 KO cells rescued with RNF25-WT-GFP and RNF25-C159/161S-GFP, treated or not with 3 µg/mL CHX, and **(D)** 3 µg/mL CHX plus 3 µM of cGAS inhibitor G140 for 3 hours. For all immunofluorescence experiments, three independent replicates were performed (N=3). Graphs representing nucleus versus cytoplasm ratio of individual cells per replicate are shown below every set of fluorescence microscopy images. Same colored dots represent a replicate, while outlined dots depict the median for each replicate. For statistical analysis, *p* values of unpaired two-sided t-tests were calculated by using the median’s replicates and compared between treatments. Size bars in fluorescence microscopy images represent 10 µm.

Next, decided to investigate if the ubiquitin E3 activity had a role in the nuclear localization of RNF25 in response to ribosome stalling. Therefore, we disrupted the RING domain in RNF25 by introducing the RNF25-GFP mutant in RNF25-KO cells which RING domain had been mutated to eliminate the interaction with its E2 enzyme RNF25^C159/161S^-GFP. Immunofluorescence analysis of the RNF25^C159/161S^-GFP mutant compared to the wild-type version revealed that RNF25^C159/161S^-GFP accumulated in the nucleus independently of CHX treatment (Figure 1C). This suggests that RNF25 ubiquitin E3 activity has a nuclear role in response to ribotoxic stress and it becomes trapped in the nucleus when this activity cannot be performed.

However, is possible that translation stress due to the lack of RNF25 ubiquitin E3 activity was the cause of the accumulation of the RNF25^C159/161S^-GFP mutant in the nucleus. Translation stress and collided ribosomes are co-activators of cGAS, an important sensor of damaged DNA in the cytoplasm that triggers an innate immunity response (*20*). Therefore, we measured the nuclear accumulation of RNF25-GFP and RNF25^C159/161S^-GFP in response to CHX after treating or not with the cGAS inhibitor G140 (Figure 1D). While cGAS inhibition suppressed the accumulation of RNF25 in the nucleus in response to CHX, it had no effect in the accumulation of the RNF25^C159/161S^ mutant. Altogether, this indicates that the accumulation of RNF25 in the nucleus is cGAS-dependent, supporting the hypothesis that the accumulation of the catalytic dead mutant in the nucleus is a direct effect of the ubiquitin E3 activity deficiency.

### Identification of RNF25 ubiquitination substrates

As our data indicates that RNF25 ubiquitin E3 activity has a nuclear role in response to translation stress and/or ribosome stalling, we decided to identify the specific RNF25 ubiquitination substrates. First, we purified the ubiquitin proteome from Parental and RNF25-KO cells stably expressing His-Ubiquitin and subsequently identified it by mass spectrometry-based proteomics after treating or not with the proteasome inhibitor MG132 (Supplementary Dataset 1). Proteins less or not ubiquitinated in RNF25-KO could be considered putative RNF25 ubiquitination substrates. However, statistical analysis did not identify proteins that were not ubiquitinated in the RNF25-KO cells in a significant manner. Moreover, this approach does not enable to distinguish direct substrates of changes in the ubiquitination status of a protein due to indirect effects.

Therefore, we applied the TULIP2 methodology which enables the identification of specific E2/E3 ubiquitination substrates in a direct manner (*21–23*). In brief, in this method, a linear fusion between an E2/E3 enzyme of interest and ubiquitin is performed. The rationale behind is that the E2/E3 will employ its attached ubiquitin moiety to modify its substrate, creating a covalent fusion between the E2/E3, ubiquitin and the substrate that can later be co-purified and identified by mass spectrometry based proteomics. As negative controls for the wild-type RNF25-TULIP2 construct, we expressed a construct of RNF25 fused to ubiquitin lacking the C-terminal diGly motif (ΔGG) and RNF25^C159/161S^ mutant which interaction with its E2 enzyme is impaired (Figure 2A).

**Figure 2.**
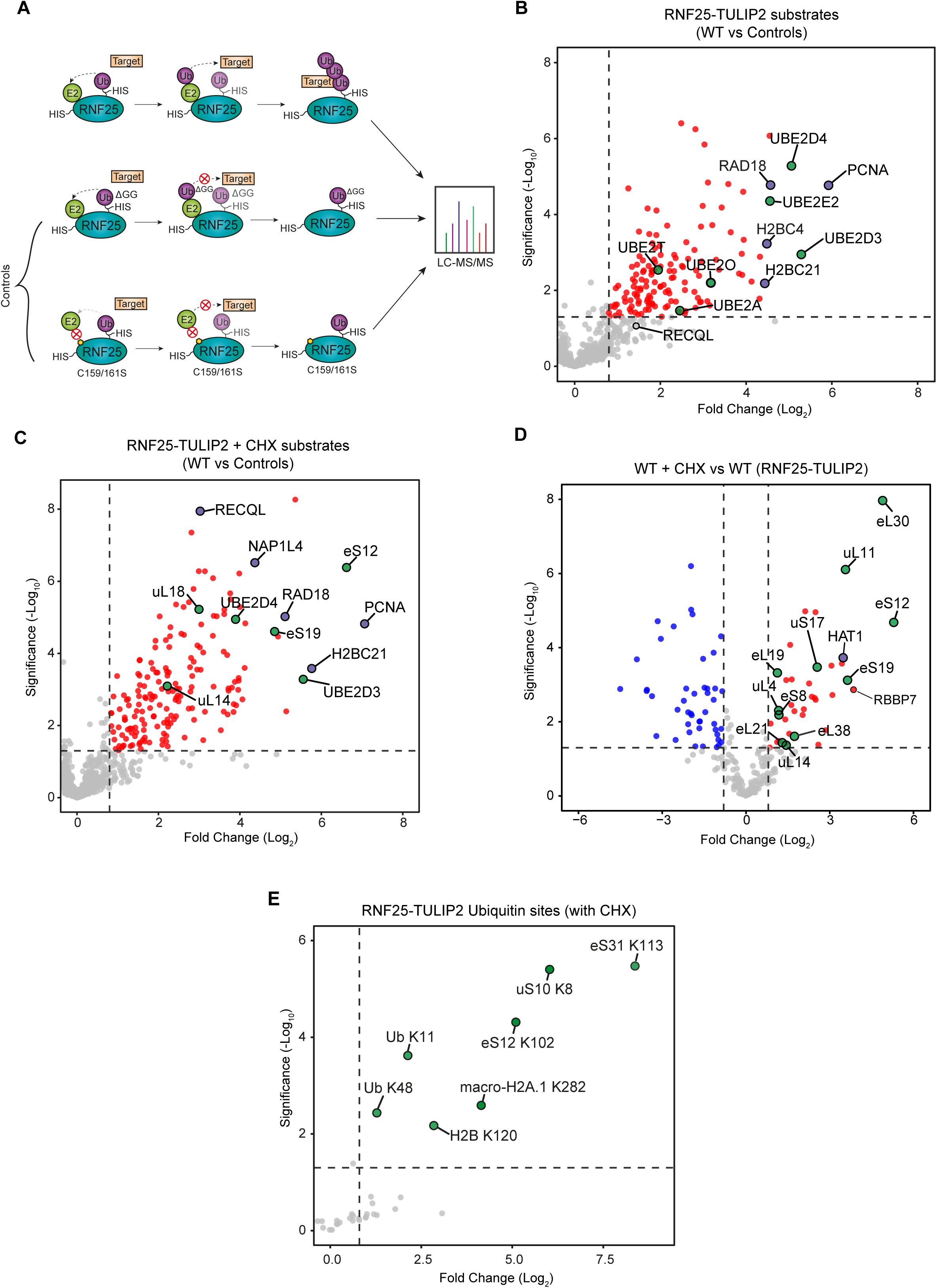
Identification of RNF25 direct ubiquitination substrates using TULIP2 methodology. **(A)** Cartoon representing RNF25-TULIP2 rationale. **(B, D)** Volcano plots depicting statistical differences between RNF25-WT-TULIP2 versus RNF25-TULIP2-ΔGG and RNF25-C159/161S-TULIP2 (named as Controls), **(C)** treated or **(B)** not with CHX, and **(D)** RNF25-WT-TULIP2 with and without CHX. Each dot represents a protein, and selected proteins are labeled. Proteins related to similar biological processes are color matched. Dashed lines mark significance threshold for *p* > 0.05 from unpaired two-tailed t-tests and fold change higher than 0.8 (log_2_). **(E)** Volcano plot showing statistical differences between GlyGlyK sites of RNF25-WT-TULIP2 and RNF25-WT-TULIP2-ΔGG under treatment with CHX. Ubiquitin chains are referred as Ub-lysine (K)-chain number, and ubiquitination lysine sites (K) on specific proteins are labeled. (F-I) Immunoblots of Parental and RNF25 KO inputs against (F) RAD18, (G) PCNA monoubiquitinated on lysine K164, (H) histone H2B monoubiquitinated on lysine K120, and (I) RECQL. Ponceau staining is shown as loading control.

We performed our RNF25-TULIP2 assays in unperturbed conditions and after treating the cells with CHX (Figure 2, Supplementary Figure 3, Supplementary Dataset 2, 3). When comparing high confidence RNF25-TULIP2 substrates, namely proteins that were enriched when comparing the wild-type TULIP2 construct with the ΔGG construct, the RNF25^C159/161S^ -TULIP2 mutant construct, and both together combined, in unperturbed conditions, the most noticeable hits were Histone H2B, PCNA, RAD18 and several Ubiquitin E2 enzymes, including UBE2D3, UBE2D4, UBE2E2, UBE2O, among others (Figure 2B). Main ubiquitin substrates identified by RNF25-TULIP2 were more highly enriched after treatment with CHX (Figure 2C-D). Additionally, other prominent CHX-enriched RNF25-TULIP2 substrates include the RECQL helicase, NAP2 and HAT1, involved in nucleosome assembly, and several ribosomal subunits involved in protein quality control. Noteworthy, ribosomal subunits involved in protein quality control have been recently described as ubiquitination substrates for RNF25 (*7*).

### UBE2D3 and UBE2D2 are the main E2 enzymes for RNF25

Many different E2 enzymes were identified as RNF25-TULIP2 substrates. However, UBE2D2, which is the E2 which has been described to act with RNF25 (*24*), was no co-purified in our RNF25-TULIP2 experiments. Nevertheless, there is the possibility that the covalent attachment of the E2s to the TULIP2 constructs was due to the binding of the TULIP2 ubiquitin moiety to the catalytic cysteine of the E2 instead of a substrate lysine. However, our acquired mass spectrometry data did not confidently identify di-Gly ubiquitin remnant motifs neither in on cysteines nor lysine corresponding to E2 enzymes.

To be able to distinguish between substrates and partner E2s to the RNF25 E3 enzyme, we performed both GFP-Trap co-immunoprecipitations of the RNF25-WT-GFP and RNF25^C159/161S^-GFP mutant expressing cells and TurboID assays both with RNF25 and RNF25^C159/161S^ TurboID-tagged constructs and subsequent analysis of the purified proteins by mass spectrometry-based proteomics (Figure 3, Supplementary Dataset 4, 5). While the GFP-Trap experiments enable the identification of stable interactions, the Turbo ID assays facilitate the detection of more transient interactions (Figure 3A-B). TurboID experiments revealed the interaction of RNF25 with a variety of E2 enzymes, including UBE2D1, UBE2D2, UBE2D3 and UBE2E1 and this interaction was lost in the RNF25^C159/161S^ mutant (Figure 3C). However, in the case of the GFP-Trap experiments, only UBE2D2 and UBE2D3 remained as interactors of RNF25 depending on E3 activity (Figure 3D). Together, all these data suggest that, while RNF25 can act with variety of E2 enzymes, UBE2D2 and UBED3 are the preferred E2s for RNF25 followed by UBE2D1 and UBE2E1. Other E2s as UBE2O, UBE2E2, UBE2T and UBE2N might be ubiquitination substrates for RNF25.

**Figure 3.**
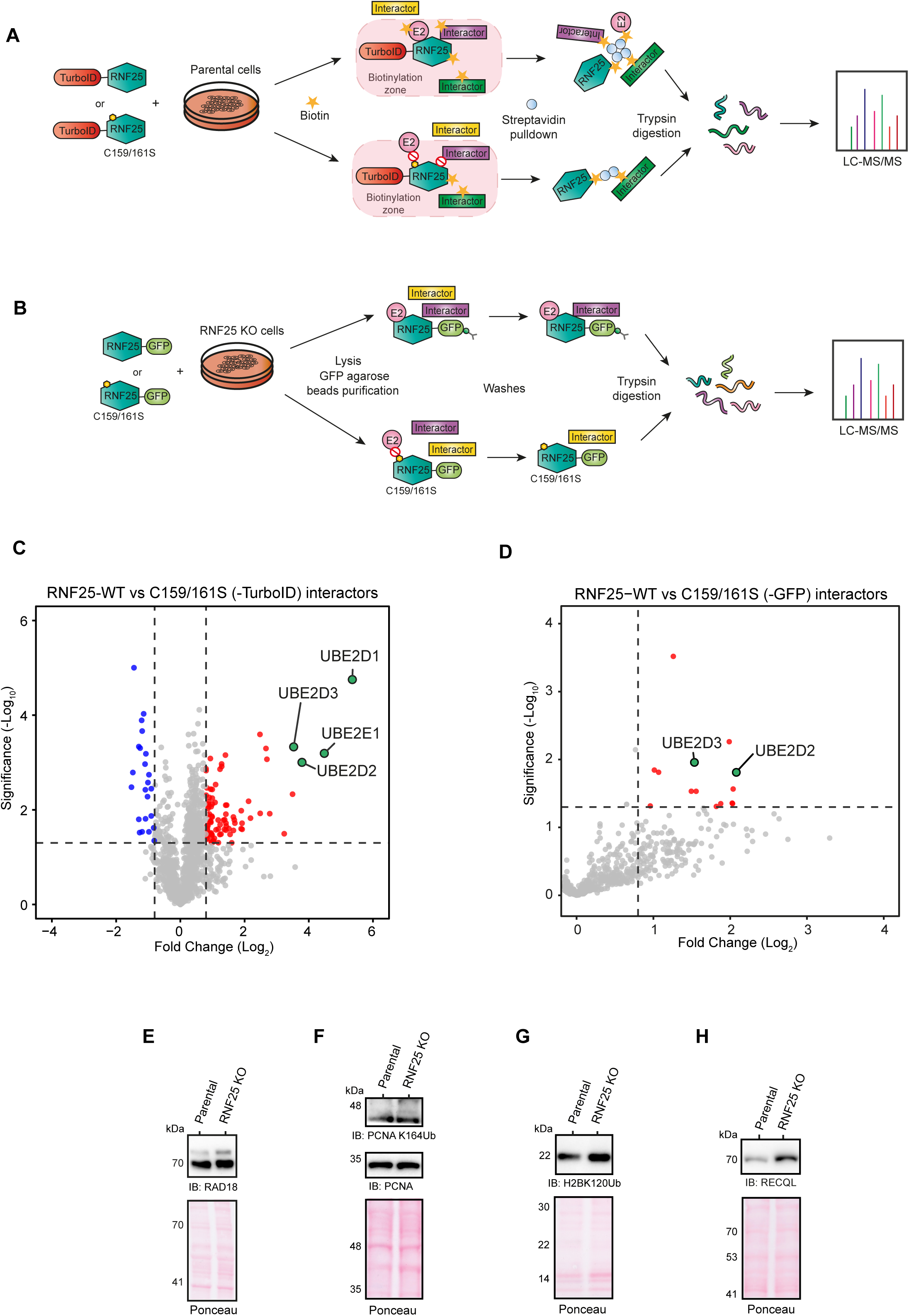
UBE2D3 and UBE2D2 are the preferred E2 ezymes for RNF25. **(A, B)** Illustration depicting TurboID and GFP co-immunoprecipitation experiments coupled to LC-MS/MS. **(C, D)** Volcano plots representing statistical differences between **(C)** RNF25-WT-TurboID and RNF25-C159/161S-TurboID, and **(D)** RNF25-WT-GFP and RNF25-C159/161S-GFP. Each dot represents a protein, and selected proteins are labeled. Ubiquitin E2 enzymes are color matched. Dashed lines mark significance threshold for *p* > 0.05 from unpaired two-tailed t-tests and fold change higher than 0.8 (log_2_).

UBE2D2 and UBE2D3 build up K48 ubiquitin chains in vitro. Additionally, UBE2D3 is able to form K11 ubiquitin chains(*25*). Consistently, analysis of ubiquitination sites in the RNF25-TULIP2 experiments showed ubiquitination on K11 and K48 sites on ubiquitin plus lysine K120 on histone H2B (Figure 2E, Supplementary Dataset 3), which favors UBE2D3 as the main E2 for RNF25.

### RNF25-TULIP2 substrates are degraded in and RNF25-dependent manner

Mixed K11/K48 ubiquitin chains signal for enhanced rapid proteasomal-mediated degradation of acceptor proteins (*26*). Recently, it has been shown that RAD18 and monoubiquitinated PCNA levels are increased at the chromatin fraction in the absence of RNF25 in response to Hydroxyurea treatment, and explained as a compensatory response to replication stress due to RNF25 deficiency(*9*). Interestingly, RAD18 and PCNA were identified as top RNF25 ubiquitination substrates in our RNF25-TULIP2 experiments already in unperturbed conditions (Figure 2B). Thus, we hypothesized that RNF25 was promoting the ubiquitination and rapid degradation of RAD18 and PCNA via K11/K48 chains. Accordingly compared to Parental cells, RAD18 and mono-ubiquitinated PCNA protein levels were increased in RNF25-KO cells (Figure 3E-F).

The most understood form of ubiquitination on PCNA consist either on mono ubiquitination, which promotes Trans-Lesion Synthesis (TLS) or K63 ubiquitin chains promoting Template Switching (TS) (*27*). To investigate if K11 and K48 ubiquitin chains are also formed on PCNA in unperturbed conditions and its dependance on RNF25, we expressed a GFP-PCNA construct in Parental and RNF25-KO cells and purified it using GFP-Trap under denaturing conditions to avoid the co-purification of PCNA interactors and analyzed it by mass spectrometry. Parental cells not expressing GFP-PCNA were used as negative control (Supplementary Figure 4, Supplementary Dataset 6 and 7). PCNA and Ubiquitin were significantly enriched compared to control in both Parental and RNF25-KO cells expressing GFP-PCNA. Furthermore, K11, K48 and K63 chains could be detected. Importantly, K11 ubiquitin chains showed higher levels in the RNF25-KO cells than in the Parental cells, indicating the K11-ubiquitinated PCNA stabilizes in the absence of RNF25.

Additionally, another RNF25-TULIP2 substrate was histone H2B ubiquitinated on lysine K120 (Figure 2B). Consistently, H2B-K120ub levels were also slightly increased in RNF25-KO cells compared to Parental cells (Figure 3G). Likewise, RECQL, another RNF25-TULIP2 substrate (Figure 2D) presented higher levels of expression in RNF25-KO cells (Figure 3H).

### RNF25 ubiquitin E3 activity promotes continuous DNA synthesis

Recently, a replication fork protective mechanism mediated by RNF168, another ubiquitin E3 enzyme, preventing toxic RAD18 accumulation has been described (*28*). Misregulation of PCNA ubiquitination causes replication problems in unperturbed conditions by the accumulation of single-stranded DNA (ssDNA) gaps left behind the replication fork (*21, 29*). Moreover, based replication dynamics, ssDNA accumulation and replication fork stability upon replication stress in Pancreatic Ductal Adenocarcinoma (PDACs) and Non-Small Cell Lung Carcinoma (NSCLC) cell lines, a role in DNA replication, yet E3 activity-independent, for RNF25 has been proposed(*8*).

Thus, we decided to investigate replication fork dynamics and the accumulation of ssDNA gaps behind the fork in Parental and RNF25-KO, non-cancer-derived RPE1 cells, rescued or not with either RNF25-WT-GFP or RNF25^C159/161S^-GFP catalytic-dead mutant by performing S1-nuclease fiber assays (*30*) (Figure 4A). In contrast to previously observed in PDACs and NSCLCs cells, our RPE1 RNF25-KO cells presented a slightly, but significant, faster replication fork progression rate on the expense of accumulation of ssDNA gaps behind the replication fork. This phenotype was corroborated in an additional RNF25-KO clone (Supplementary Figure 5A). Importantly, expression of RNF25-WT-GFP, but not RNF25^C159/161S^-GFP, rescued this phenotype. Moreover, we performed clonogenic assays in response to treatment with WEE1 inhibitor, a drug RNF25-KO cells are higly sensitive to (*9*) (Figure 4B). In a similar manner, expression of RNF25-WT-GFP, but not RNF25^C159/161S^-GFP, rescued the sensitivity of RNF25-KO cells to WEE1 inhibition. Likewise, the equivalent, previously described (*9*) RNF25^C135/138S^-GFP E3-deficient mutant was also sensitive to WEE1 inhibition (Supplementary Figure 5B)Together, our results indicate that, in contrast to recently described, the DNA replication protective function of RNF25 is dependent on its ubiquitin ligase function.

**Figure 4.**
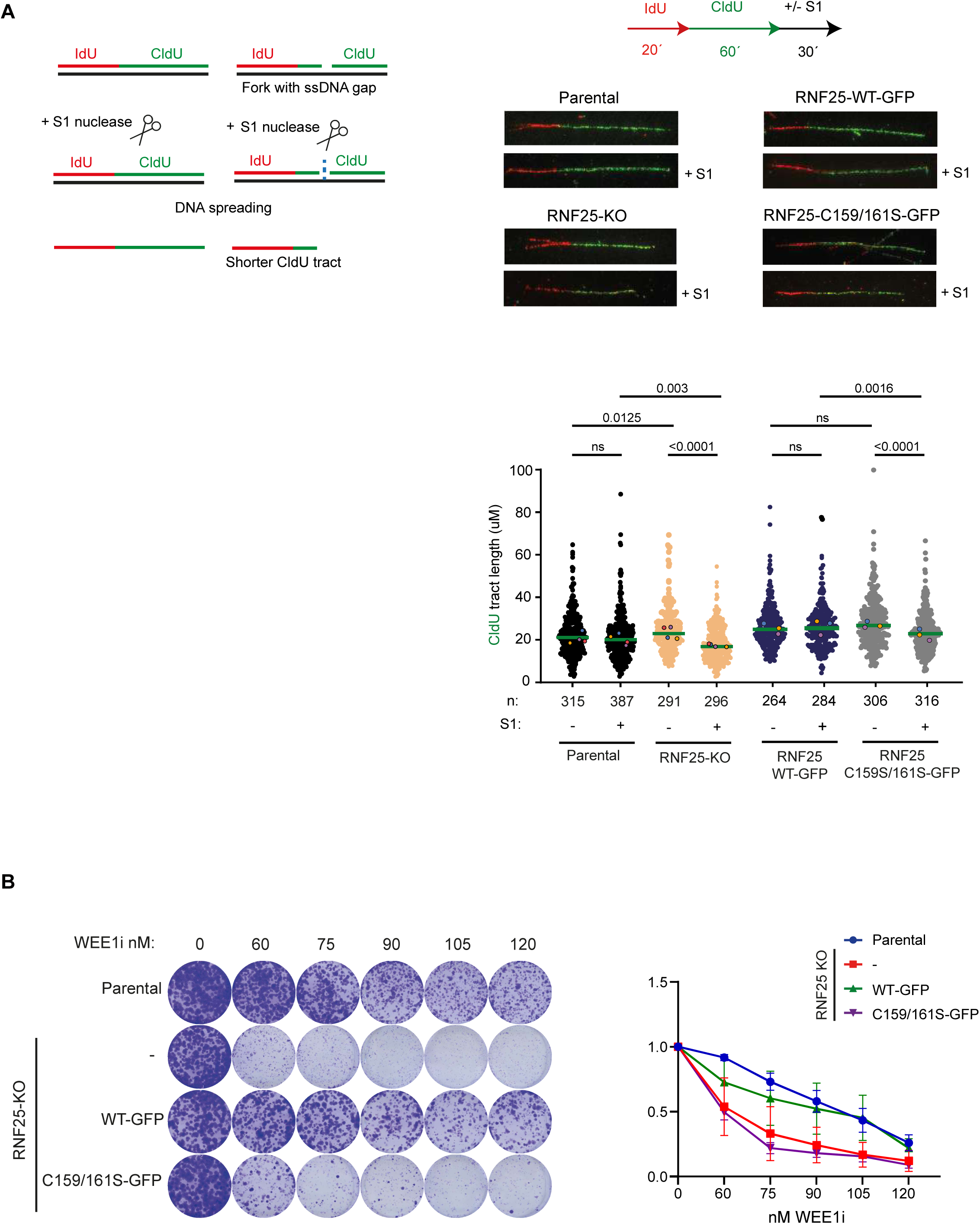
RNF25 ubiquitin E3 activity facilitates continuous DNA synthesis. **(A)** Top left: Drawing of S1 nuclease DNA fiber assay protocol. Forks are labeled with IdU and CldU and subsequently treated with S1 nuclease to process ssDNA gaps. Top right: Summary of the S1 nuclease DNA fiber assay and fluorescence microscopy images representing DNA labeled fibers for Parental and RNF25 KO cells rescued or not with RNF25-WT-GFP and RNF25-C159/161S-GFP. Down right: Graphs depicting CldU tract lengths of each cell line with and without S1 nuclease treatment. Each dot represents one fiber, and colored dots indicate the median of each independent experiment. The green bar represents the median for all three replicates and *n* indicates the total fiber count per cell line and treatment. *p* values were calculated using the two-sided Mann-Whitney test. **(B)** Clonogenic survival assay of Parental and RNF25 KO cells rescued or not with RNF25-WT-GFP or RNF25-C159/161S-GFP, under treatment with WEE1 inhibitor for 24 hours and 7-days recovery. Average and standard deviation of three independent experiments with three technical repeats are shown (N = 3).

Additionally, we performed the S1 nuclease fiber assay in the presence of CHX (Supplementary Figure 5C). However, CHX treatment inhibited DNA replication, not enabling to observe differences between Parental and RNF25-KO cells.

### Deficiencies in RNF25 ubiquitin E3 activity causes Transcription-Replication Conflicts (TRCs)

In order to obtain mechanistical insight on the cause of accumulation of ssDNA gaps behind the replication fork in RNF25-KO cells, we decided to investigate proteome dynamics at replication forks by performing iPOND experiments coupled to mass spectrometry-based proteomics in Parental and RNF25-KO cells in the presence or absence of CHX (Figures 5A-B, Supplementary Figure 6, Supplementary Dataset 8). CHX treatment mainly causes a defect of histone supply to the replisome both in Paretal and RNF25-KO cells (Supplementary Figure 6C-D). Interestingly, proteins related to the surge of TRCs were recruited to replication forks in RNF25-KO cells compared to Parental cells. These, among others, included proteins involved in mRNA processing, including splicing factors and RNA helicases that resolve R-loops formation, such as DDX17 and DDX5 (*31–33*). Additionally, TRIM28, UBA2 and SUMO2, which also protect replication forks upon TRCs (*34*) or BCCIP (*35*), were also recruited to the replication forks in the absence of RNF25. Moreover, several proteins which limit the degradation of nascent DNA and DNA end resection as BRCC36 (*36–38*), or the members of the KU complex KU70/KU80 (*39*) were also enriched (Figure 5B). Furthermore, RECQL, which promotes the restart of reversed forks (*40*), was among the most highly enriched proteins at the replication fork in the absence on RNF25, and TRCs are an important factor for the induction of reversed forks (*41*). Strikingly, RECQL was found as one of the most relevant direct ubiquitination substrates for RNF25 (Figure 2C).

**Figure 5.**
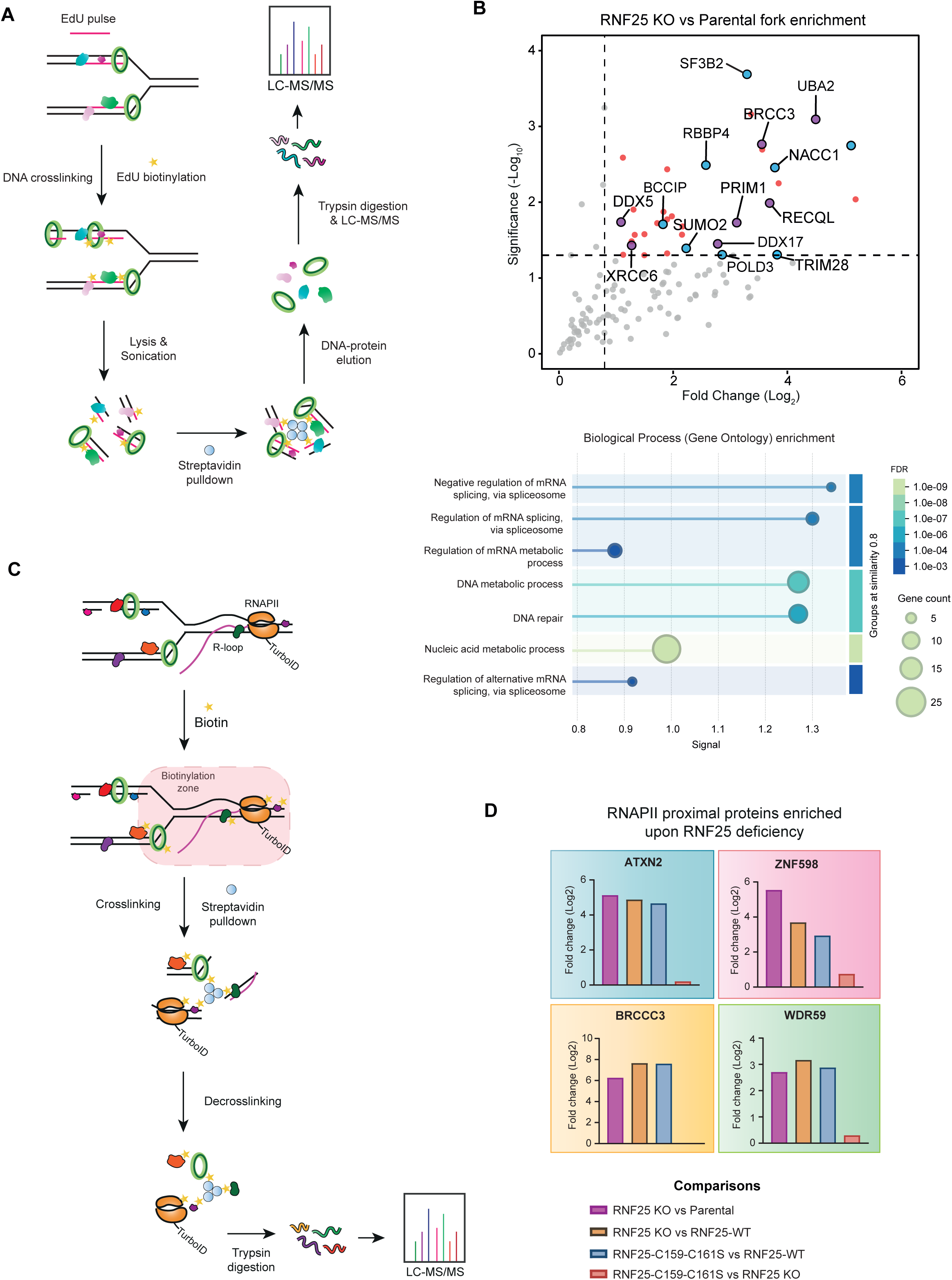
Loss of RNF25 results in enrichment of Transcription-Replication Conflicts (TRCs) markers on replication forks. **(A)** Scheme representing isolation of Proteins On Nascent DNA (iPOND) coupled to LC-MS/MS experiment. **(B)** Left: Volcano plot depicting statistical differences between replication fork enrichment of RNF25 KO and Parental. Each dot represents a protein, and selected proteins are labeled. Proteins related to similar biological processes are color matched. Dashed lines mark significance threshold for *p* > 0.05 from unpaired two-tailed t-tests and fold change higher than 0.8 (log_2_). Right: STRING gene ontology enrichment of significant replication fork-associated proteins in RNF25 KO. **(C)** Cartoon depicting POLR2I-TurboID strategy. **(D)** Bar plot representing statistical differences for ATXN2, ZNF598, BRCC3 and WDR59 proteins between RNF25 mutants (RNF25 KO and RNF25-C159/161S-GFP) and WT (Parental and RNF25-WT-GFP) expressing POLR2I-TurboID constructs. Each colored bar shows the comparisons between samples.

Nevertheless, TRCs involve not only DNA replication but also transcription. Therefore, we decided to perform a proximal proteomic characterization of RNA polymerase II using TurboID(*42*). For this purpose, we stable-inducible expressed POLR2I-TurboID in Parental and RNF25-KO cells (Supplementary Figure 7A-B). However, POLR2I-TurboID was more highly expressed in RNF25-KO than in Parental Cells, which might be difficult for the interpretation of the results. As we were unable to achieve similar expression levels of POLR2I-TurboID in Parental and RNF25-KO cells, we rescued or not again POLR2I-TurboID RNF25-KO cells with either RNF25-GFP or RNF25^C159/161S^-GFP. First, we compared the RNA polymerase II proximal proteome in the different cell lines (Figure 5C-D, Supplementary Figure 7C, Supplementary Dataset 9) and investigated proteins that become in proximity of RNA polymerase II in the absence of RNF25 ubiquitin E3 activity, that is, proteins that were enriched both in the RNF25-KO and RNF25^C159/161S^-GFP rescued cells compared to both Parental and RNF25-GFP rescued cells. Twelve different proteins were identified to fulfill these conditions (Suplementary Figure 7D, Figure 5D), including BRCC3, which is also recruited to replication forks in the absence of RNF25 (Figure 5B). Interestingly, ZNF598, which is synthetic lethal with RNF25 in response to RNA damage(*7*) interacted with RNA polymerase in absence of RNF25. ZNF280C, another protein which localizes at and protects stalled replication forks, (*43*) also becomes in proximity to RNA polymerase II in the absence of RNF25. Finally, ATXN2, a RNA-binding protein involved in R-loop suppression(*44*), becomes in proximity of RNA polymerase II in the absence of RNF25 ubiquitin E3 activity.

Taken together, all these data strongly confirms that deficiency in RNF25 induces the occurrence of TRCs. TRCs have been shown to promote the accumulation of ssDNA gaps behind the replication fork due to repriming ahead of R-loops (*41*). Consistently, we observed increased levels of primase and R-loop processing proteins at replication forks (Figures 5B and 5D, Supplementary Datasets 8, 9). Therefore, we hypothesized that suppressing R-loop formation should rescue the replicative ssDNA gap formation phenotype in RNF25-KO cells. To confirm our hypothesis, we transiently over expressed or not RNAseH1 in Parental and RNF25-KO cells and performed the S1-nuclease fiber assay (Figure 6A-B). As predicted, RNAseH1 overexpression suppressed the accumulation of ssDNA gaps behind replication forks in RNF25-KO mutants.

**Figure 6.**
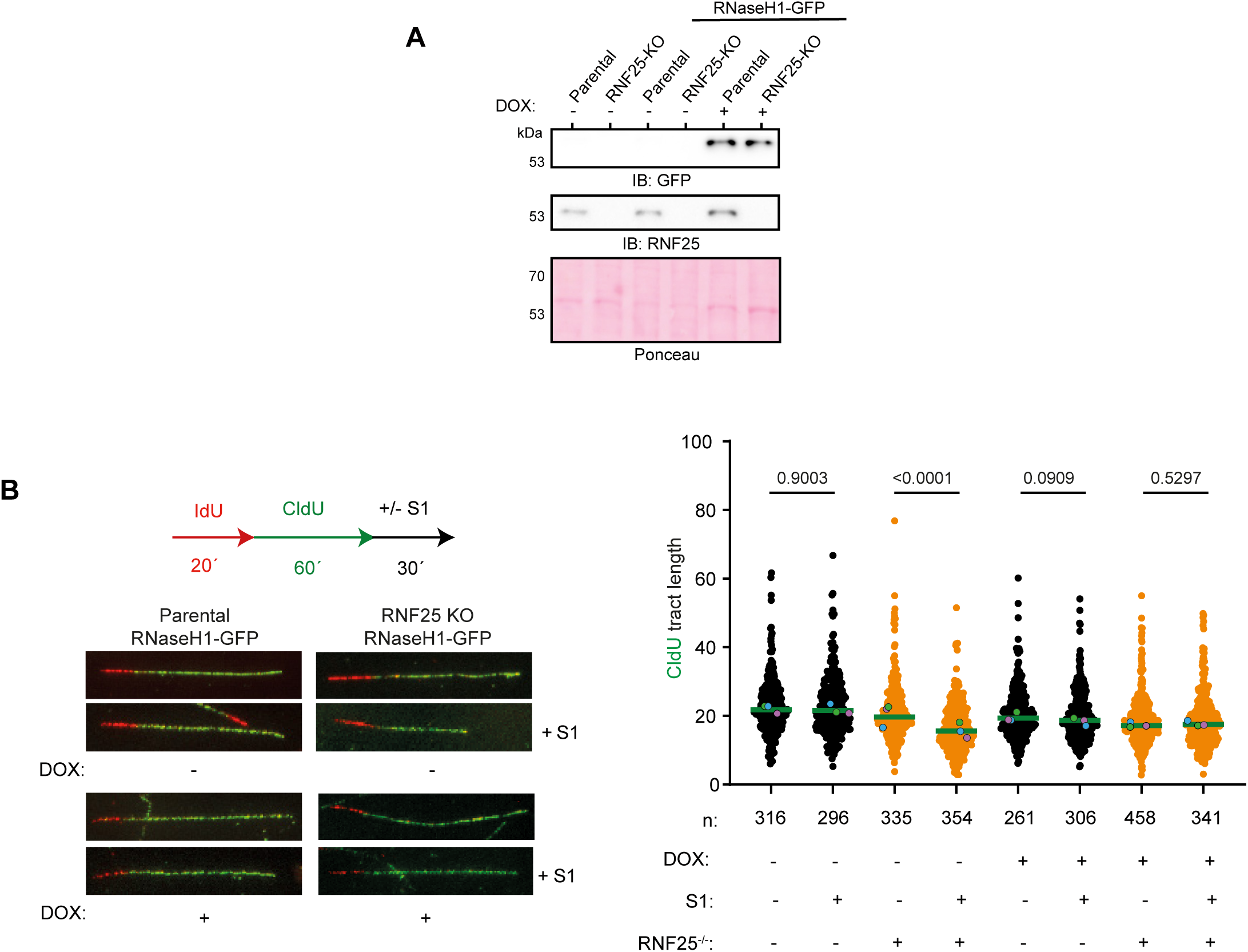
RNaseH1 rescues the replication fork progression in RNF25 mutant cells. **(A)** Immunoblotting of Parental and RNF25 KO cells expressing or not RNaseH1-GFP. Induction of the construct was performed with doxycycline at 5 µg/mL for 48 hours. **(B)** Up: Fluorescence microscopy images representing labeled IdU and CldU fibers of S1 nuclease DNA fiber assay from Parental and RNF25 KO cells expressing or not RNaseH1-GFP. Down: Graphs depicting CldU tract lengths of each cell line with and without S1 nuclease treatment. Each dot represents one fiber, and colored dots indicate the median of each independent experiment. The green bar represents the median for all three replicates and *n* indicates the total fiber count per cell line and treatment. *p* values were calculated using the two-sided Mann-Whitney test.

### RNF25 E3 stability preserves replication fork stability by coordinating Translation, Transcription and DNA replication

Upon translational stress, RNF25 has been shown to prevent the hyperactivation of GCN2-dependent Integrated Stress Response (ISR) via the ubiquitination of the ribosomal subunits eS31in lysine K113 and uL11, which was indirectly determined by differences in diGly proteomics between control and RNF25-KO HAP1 cells (*7*). Our RNF25-TULIP2 data confirmed the ubiquitination of both the eS31 in lysine K133 and uL11 in a direct manner plus other ribosomal proteins as eS12, eL30, eS19, uS17, eL38, uL14, eL21, eS8 and uL4 which are ubiquitinated by RNF25 in response to translational stress caused by CHX treatment (Figure 2D, Supplementary Dataset 2, 3).

Moreover, activation of the ISR promoted the translocation of RNF25 to the nucleus in a cGAS-dependent manner (Figure 1). In the nucleus, RNF25 has been previously shown to localize at DNA replication sites by SIRF experiments(*8*). At the replisome, RNF25-dependent branched K11/K48 ubiquitin chains would promote the rapid clearance of proteins by the proteasome at different levels to solve TRCs.

We propose a model in which degradation of H2B-K120Ub, a mark that promotes transcription elongation, is a mechanism to reset transcription. This transcriptional reset would suppress R-loop formation at the fork. Moreover, H2B-K120Ub is a mark that promotes lesion bypass during replication(*45*), a mechanism also dependent on RAD18 and PCNA-Ub (*27*) which would be also cleared from the replication fork by RNF25. Clearance of H2B-K120Ub, RAD18 and PCNA-Ub would shift DNA Damage Tolerance towards a fork reversal mechanism to stabilize stressed replication forks. Furthermore, clearance of RECQL would prevent the premature dissolution of the reversed fork promoting replication fork stability. On the other hand, in an RNF25-deficient scenario, higher levels of TRCs would require the recruitment to the fork of several factors that stabilize the replication fork in response to TRCs, including RECQL, which could resolve the reversed fork to enable re-priming downstream the TRC as previously described (*46*) leaving a gap behind the fork (Figure 7). In a similar manner, yet mechanistically distinct, deficiencies in other factors, such as BRCA1 and PCNA ubiquitination promote fork degradation upon stalling (*29, 47*), but present discontinuous DNA synthesis in unperturbed conditions(*21*).

**Figure 7.**
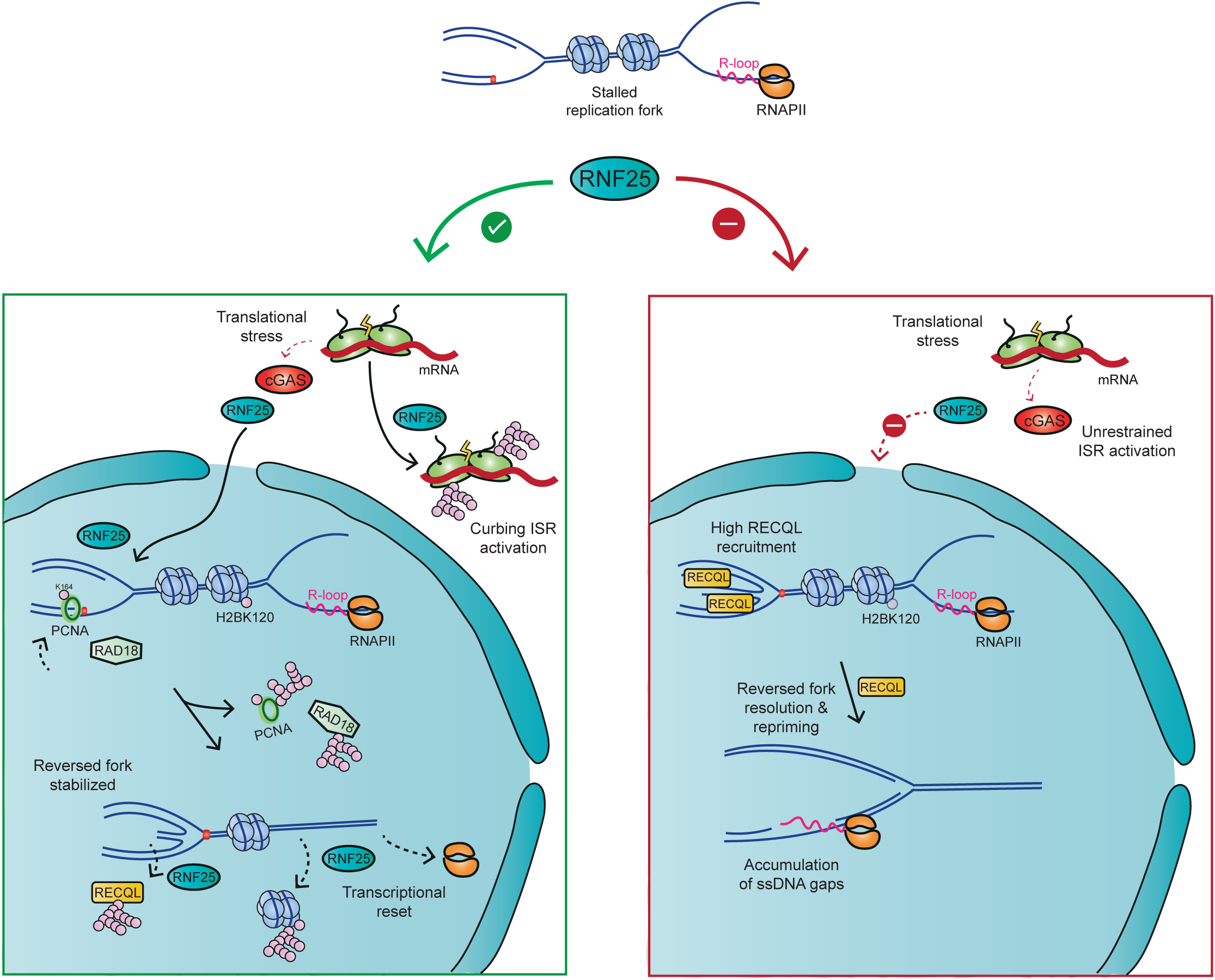
Model. RNF25 coordinates replication, transcription and translation through its E3 ligase activity. Proposed model for RNF25 role coordinating replication, transcription and translation in two different scenarios. Upon replication fork stalling, RNF25 translocate to the nucleus in a cGAS dependent manner to mark H2B-K120, RAD18 and PCNA for degradation, stabilizing the stressed forks by fork reversal activation. Clearance of RECQL would promote fork stability by preventing the premature dissolution of the reversed fork. However, in RNF25-deficient cells, RECQL would accumulate in the replication fork by sensing higher levels of TRCs and would resolve the reversed fork structure to enable re-priming downstream the TRC, leaving a gap behind the fork.

Previous observations proposed that, in a situation of fork stalling due to high hydroxyurea treatment, RNF25 and REV7 interact and participate in the same reversed fork stabilization pathway (*8*). However, we did not find REV7 in any of our different mass spectrometry-based proteomics analysis, including GFP-Trap coimmunoprecipitation, RNF25-TurboID, RNF25-TULIP2 or iPONDs. Nevertheless, our model is fully compatible with these previous observations. Lack of REV7 or RNF25 would promote the dissolution of the same reversed forks by distinct mechanisms. Namely, lack of REV7 would degrade reversed forks because the nucleolytic action of MRE11 and CtIP, and lack of RNF25 would restore the reversed fork by RECQL exposing the stalled fork to degradation by MRE11 and CtIP. Here, repriming would not be possible due to the hydroxyurea treatment, explaining the epistatic effect.

In our model, deletion of RECQL should suppress the resolution of the reversed forks in RNF25-KO cells, disabling the re-priming tolerance pathway to TRCs. To test this hypothesis in an unbiased manner, we performed a genome-wide synthetic viability/lethality CRISPR-Cas9-based screen in unperturbed conditions. We infected Parental and RNF25-KO cells with the Brunello Human CRISPR Knockout pooled one-vector system library consisting of 76,441 gRNAs targeting 19,114 human genes (*48, 49*). After 21 doublings, we sequenced the gRNAs and obtained the results from the genome-wide CRISPR-Cas9 knockout screen (Figure 8A-B, Supplemental Dataset 10). As expected, deletion of WEE1 and related genes such as CDK1 or CNTD1 were deleterious to RNF25-KO cells. Supporting our hypothesis, RECQL knockout provided a significant growth disadvantage in RNF25-KO cells but not in Parental cells. ZNF598 knockout, which is synthetic lethal with RNF25 knockout upon azacitidine treatment(*7*),was also detrimental to RNF25-KO cells in unperturbed conditions.

**Figure 8.**
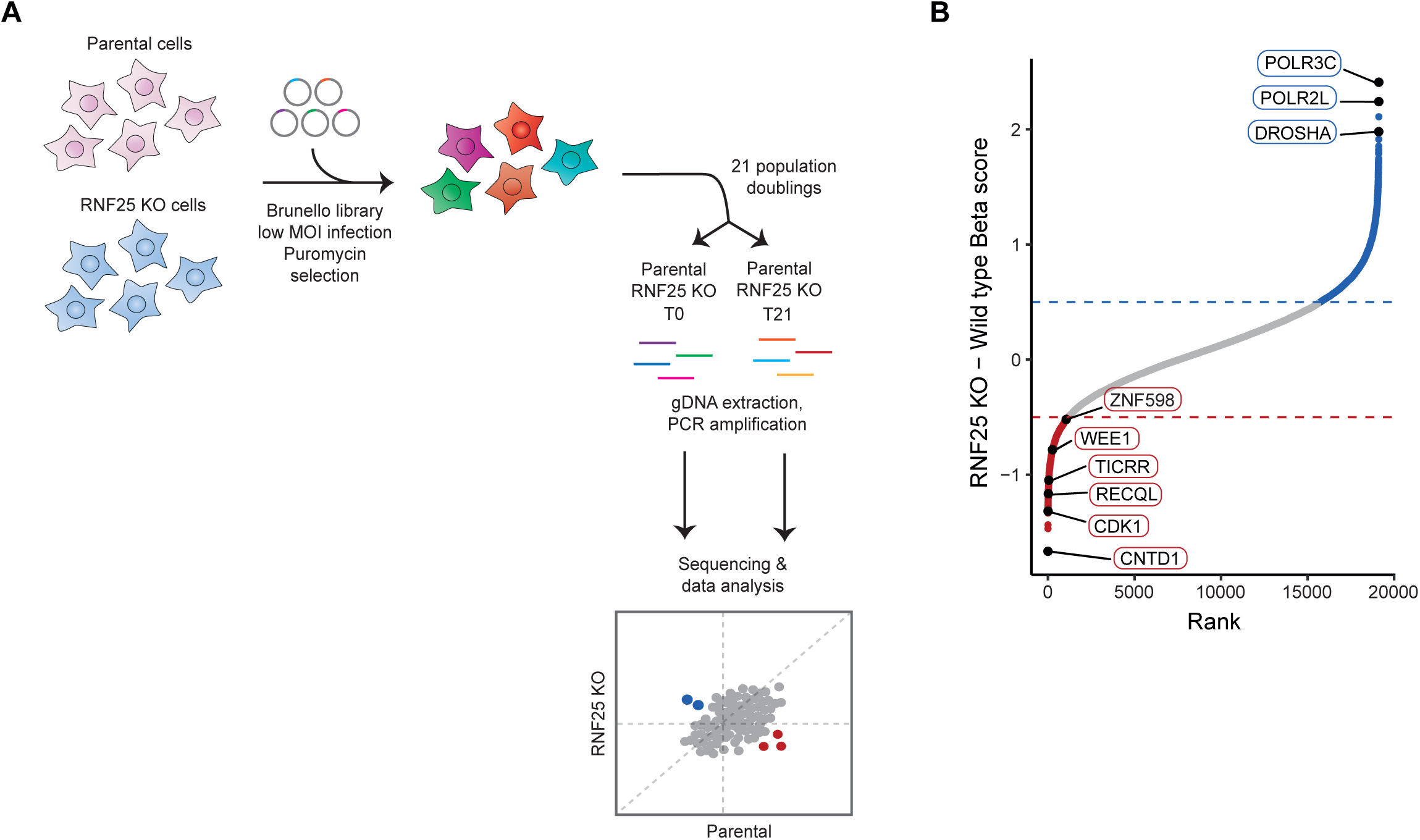
CRISPR screen identifies gene candidates that are synthetically lethal or viable under a RNF25 depletion. **(A)** Synthetic lethality CRISPR/Cas9 screening experimental design. **(B)** Gene ranks of the differential beta score between RNF25 KO and Parental cells, where each dot represents a gene. Blue or red dots have a differential beta score higher than 0.5 or less than 0.5, respectively. Highlighted dots are synthetic lethality and viability candidate genes for RNF25.

## DISCUSSION

RNF25 has recently emerged as a key ubiquitin E3 enzyme in cell tolerance against different types of RNA damage causing ribosome stalling (*7, 10, 11*). In these scenarios, RNF25 mediates the degradation of different translation factors at the ribosome. Using TULIP2 methodology, we found that, additionally to these ribosomal factors, nuclear proteins involved in DNA replication and transcription regulation are also ubiquitinated by RNF25. Importantly, upon translational stress, we observed that RNF25 accumulates in the nucleus depending on the ISR mediated by cGAS where it promotes the degradation of proteins involved in replication fork stability and transcription. This function would serve to coordinate translation with transcription and DNA replication.

On one side, translation and replication need to be highly coordinated as newly synthesized DNA requires a high supply of histones to conform the new chromatin. In situations of translational stress, halting DNA replication and reversing replication forks would be physiologically beneficial to stabilize forks avoiding their collapse. At the level of transcription, transcriptional reset would be beneficial to avoid the occurrence of TRCs, specially at highly transcribed genes, which in human cells are marked by histone H2B ubiquitination at lysine K120 (*50*). Our data shows the RNF25 promotes rapid clearance by ubiquitination of proteins which activity would be detrimental in these scenarios.

A role for RNF25 in the stabilization of replication forks was already recently described (*9*), and our data strongly support these observations. However, some discrepancies between the previously published work and our results have arisen. First, while in the previous work of the Vaziri laboratory they showed that RNF25 knockout had a slower replication rate compared to the Parentals, we did not observe such phenotype. In contrast, we observe a slightly higher replication rate on RNF25-KO cells compared to Parental cells on the expense of ssDNA gap formation behind the gap. This contrast can be explained by the difference in the cell lines used as model in each study. While we used non-cancer-derived RPE1 cells, in the Varizi lab work, they employed cells derived from PDACs and NSCLC. PDACs and NSCLC present high levels of replication stress and TRCs, thus, the deletion of RNF25 would be much more deleterious in these cells (*51, 52*).

Another important discrepancy between our work and the previous work from the Vaziri lab is the relevance of the ubiquitin E3 activity for the replicative function of RNF25. In their data, deletion of the RING domain in RNF25 phenocopied the RNF25 knockout in regard of its replicative function. However, introducing the C135/138S mutation to disrupt the binding to the E2, caused no effect, and concluded that the E3 activity was not required. In contrast, we saw that introducing the equivalent C159/161S mutation to disrupt the interaction with the E2s, had the same consequences that RNF25 knockout for ssDNA gap formation during replication. Importantly, the only stable interactors of RNF25 lost with the C159/161S mutation were UBE2D2 and UBE2D3.

Finally, while the Vaziri lab linked, among others, the replicative defect phenotype of RNF25 loss to an interaction with REV7, we were unable to identify REV7 in any of our mass spectrometry-based proteomics data. We have performed many different approaches, TULIP2 for the identification of ubiquitination substrates, GFP-Trap coimmunoprecipitation and TurboID to map the proximal interactome including transient interactions. This discrepancy can be explained by differences in the stringency of the co-immunopurification conditions.

Nevertheless, conclusions both from our work and from the Vaziri lab are compatible. The orthogonally validated mass spectrometry approaches that we applied in this work provided mechanistical insight into the role in reversed fork protection earlier proposed by the Vaziri lab. However, our data supports an alternative model in which RNF25 would be promoting the rapid clearance of RECQL from the reversed fork to prevent is premature resolution and not to recruit REV7 to prevent their degradation by nucleases as MRE11 or CtIP.

## MATERIALS AND METHODS

### Generation of constructs

RNF25 cDNA without STOP codon was cloned into pDNOR223 by Gateway® cloning BP reaction. Site-directed mutagenesis over pDONR223-RNF25 was performed to achieve a catalytic dead mutant with C159/C161S mutations. GFP-PCNA cDNA was gifted by Caren Norden (Addgene, #105937) and cloned into pDONR207. Lentiviral constructs were generated by Gateway® cloning LR reaction between donor plasmids and acceptors TULIP2(*22*), TurboID(*42, 53*) or pLX303 (Addgene, #25897)(*54*). For the RNF25-GFP constructs, the LR reactions were performed with pLVU-GFP, gift from Lars Ittner (Addgene, #24177)(*55*). The RNaseH1 inducible plasmid (pTSB27_NA85-pLPT-M27-RNaseH1-EGFP) was kindly provided by Andrés Aguilera and Emilia Herrera-Moyano. All primers and plasmids are listed in Supplementary Table 1 and 2.

### Cell culture

HEK 293T and RPE1-*TP53^-/-^PAC^-/-^* (Parental) cells were cultured in Dulbecco’s modified Eagle’s medium (DMEM) supplemented with 10 % Fetal Bovine Serum (FBS) and Penicillin (100 U/mL)/ Streptomycin (100 µg/mL) (Pen/Strep) at 37°C and 5% CO_2_ unless specifically specified. Cells were regularly tested for mycoplasma contamination.

### Lentivirus production

HEK 293T cells were seeded at 30% confluence in a T175 flask containing 17 mL of DMEM + FBS. After 24 hours, the transfection mixture was prepared by combining lentiviral packaging plasmids: 7.5 µg pMD2.G (Addgene, #12259), 11.4 µg pMDLg-RRE (Addgene, #12251), 5.4 µg pRSV-REV (Addgene, #12253) and 13.7 µg of generated constructs (listed in Supplementary Table 2) with 114 µL of Polyethylenimine (PEI, 1 mg/mL) in 2 mL 150 mM NaCl. The mixture was vortexed and incubated 15 min at room temperature (RT) before adding to the HEK 293T cells. Next day, culture medium was refreshed with DMEM + FBS. 72 hours post-transfection, lentiviral suspension was collected and filtered through a 0.45 µm syringe filter (SLHVM33RS, Millex^®^-HV) and kept at -20 °C for further use.

### Target gene knock-out cell line generation

RPE1-*TP53^-/-^PAC^-/-^* (Parental) cells transfection mixture was prepared by combining 150 mM NaCl, 24 µg pSpCas9(BB)-2A-GFP (PX458) (Addgene, #48138) plasmid containing sgRNA targeting RNF25, and 60 µL of PEI. The mixture was vortexed and incubated 15 min at RT before adding to the cells. After 48 h, GFP positive cells were sorted, and single clones were generated by seeding cells in low density in 96-well plates (1 cell/well). After clones expanded, RNF25 depletion was validated through immunoblotting. sgRNAs sequences are listed in Supplementary Table 1.

### Generation of stable cell lines

Parental or RPE1-*TP53^-/-^PAC^-/-^ RNF25^-/-^*(*RNF25^-/-^*) cells were seeded at 15% confluence in 15 cm plates with DMEM + FBS. Next day, cell culture medium was replaced with lentivirus-containing medium RNF25-TULIP2, RNF25-TurboID, RNF25-GFP or RNaseH1-GFP constructs, along with 8 µg/mL polybrene. After 24 hours, medium was refreshed with DMEM + FBS/Pen/Strep. 72 hours post-infection, RNF25-TULIP2, RNF25-TurboID and RNaseH1-GFP positive cells were selected with 3 µg/mL puromycin. RNF25-GFP positive cells were selected with 5 µg/mL blasticidin and sorted in a BD FACSAria™ IIIu Cell Sorter based on GFP intensity.

### Electrophoresis and immunoblotting

Input samples were lysed in 200-400 μl of SNTBS buffer [2% SDS, 1% NP-40, 50 mM Tris-HCl (pH 7.5), and 150 mM NaCl] and boiled at 100°C. Samples were separated on Novex 4 to 12% gradient gels (Thermo Fisher Scientific) using NuPAGE MOPS SDS running buffer [50 mM MOPS, 50 mM Tris base, 0.1% SDS, and 1 mM EDTA (pH 7.7)] or in-house casted gels of different Acrylamide percentages. Proteins were transferred onto Amersham Protran Premium 0.45 NC nitrocellulose blotting membranes (GE Healthcare) using a Bolt Mini-Gel system (Thermo Fisher Scientific), which was used for both the gel electrophoresis and protein transfer to the membrane according to vendor’s instructions. Membranes were stained with Ponceau-S (Sigma-Aldrich) to determine the total amount of protein loaded. Next, membranes were blocked with blocking solution (8% milk, 0.05% Tween-20 in PBS) for 1 hour before primary antibody incubation. Primary antibodies were incubated overnight (o/n) at 4°C and secondary antibodies for 1-2 hours at RT. Chemiluminescence reaction was initiated with Western Bright Quantum Western blotting detection kit (Advansta-Isogen) and measured in an ImageQuant 800 (Cytiva, Malborough, MA, USA) system. Antibodies are listed in Supplementary Table 3 and 4.

### GFP-Trap Co-Immunoprecipitation

Co-Immunoprecipitation of RNF25-GFP was performed as previously described(*21*). Confluent 15 cm dishes of *RNF25^-/-^*cells rescued with GFP-tagged RNF25 wild type (WT) or C159/161S mutant were scrapped, washed in ice-cold PBS and lysed in 1 mL of ice-cold lysis buffer [20 mM Tris-HCl (pH 7.5), 150 mM NaCl, 1 mM MgCl_2_, 0.5% Triton X-100, EDTA-free protease inhibitors (Roche)]. Subsequently, protein concentration was determined by BiCinchoninic Acid (BCA) Protein Assay Reagent (Thermo Scientific). After equalization, lysates were vortexed and incubated for 1 hour at 4 °C while rotating with 500 Units of Benzonase (Millipore). Afterwards, samples were centrifuged for 1 hour at 20,000 x g at 4°C. As inputs, 20 µL of supernatants were saved per sample and the rest was incubated with 25 µL of GFP-Trap beads slurry (Chromotek) for 90 min at 4°C while rotating. Then, beads were washed three times with wash buffer [20 mM Tris-HCl (pH 7.5), 150 mM NaCl, 1 mM MgCl_2_, EDTA-free protease inhibitors (Roche)]. 10% of the beads were resuspended in LDS sample buffer 1X for immunoblotting procedures. The remaining beads were washed five times with 50 mM ammonium bicarbonate (ABC).

### Purification of TULIP2 conjugates

Following the TULIP2 methodology^1^, five 15 cm plates of RNF25-TULIP2 cells were grown to 80% confluence, and expression of TULIP2 constructs was induced with 1 µg/mL doxycycline. 24 h after doxycycline induction, cells were treated or not with 3 µg/mL Cycloheximide (CHX, Sigma-Aldrich) for 3 hours or with the proteosome inhibitor 10 µM MG-132 (Sigma-Aldrich, C2211) for 5 hours.

After treatment, cells were washed twice and scraped. Protocol was followed as indicated^2^. Cells were pelleted in 5 mL ice-cold PBS, and a small fraction of the pellet was taken as input and lysed in SNTBS buffer. After additional centrifugation, cells were lysed in Guanidinium buffer [6 M guanidine-HCl, 0.1 M Sodium Phosphate, 10 mM Tris-HCl (pH 7.8)] and snap frozen. After thawing, lysates were homogenized at RT by sonicating two times at 80% amplitude and 5 s using a tip sonicator (Branson 450 Ultrasonic Cell Disruptor, Emerson, Missouri, USA). Protein concentration was determined by BCA Protein Assay Reagent (Thermo Scientific). After equalization, lysates were supplemented with 5 mM β-mercaptoethanol and 50 mM Imidazole (pH 8.0). 100 µL of nickel-nitrilotriacetic acid-agarose (Ni-NTA) beads (QIAGEN), were equilibrated with Guanidinium buffer supplemented with 5 mM β-mercaptoethanol and 50 mM Imidazole (pH 8.0). Equilibrated Ni-NTA beads were added to the cell lysates and incubated o/n at 4°C under rotation.

After lysate-beads incubation, Ni-NTA beads were transferred with Wash Buffer 1 [6 M Guanidine-HCl, 0.1 M Sodium Phosphate, 10 mM Tris-HCl, 10 mM Imidazole, 5 mM β-mercaptoethanol, 0.2 % Triton X-100 (pH 7.8)] to an Eppendorf LoBind tube (Eppendorf). Then, beads were washed with Wash buffer 2 [8 M Urea, 0.1 M Sodium Phosphate, 10 mM Tris-HCl, 10 mM imidazole, 5mM β-mercaptoethanol (pH 8)] and transferred to a new LoBind tube with Wash buffer 3 [8 M urea, 0.1 M Sodium Phosphate, 10 mM Tris-HCl, 10 mM imidazole, 5 mM β-mercaptoethanol (pH 6.3)]. Ultimately, beads were washed twice with Wash buffer 4 [8 M urea, 0.1M Sodium Phosphate, 10 mM Tris-HCl, 5 mM β-mercaptoethanol (pH 6.3)]. After last wash, Ni-NTA beads were resuspended in 100 µL of Wash Buffer 5 [8 M urea, 0.1 M NaH2PO4/Na2HPO4, 0.01 M Tris-HCl (pH 7)] and 10% of the sample was taken as pull-down for immunoblotting. The rest of the sample was prepared for mass spectrometry.

### His10-Ubiquitin pull-down

Parental or *RNF25^-/-^* cells expressing His10-Ubiquitin were treated or not with 10 µM MG-132 (Sigma Aldrich) for 5 hours. Cells were then collected by scraping and centrifuged at 500 g for 5 min at 4 °C. Afterwards, pellets were resuspended in ice-cold PBS, and a small fraction was taken as inputs and lysed in SNTBS buffer. The remaining cells were centrifuged and were lysed in 6 mL of Guanidinium buffer [6 M guanidine-HCl, 0.1 M Sodium Phosphate, 10 mM Tris-HCl (pH 7.8)]. Lysates were sonicated twice for 5 s at 80% power in a tip sonicator (Branson 450 Ultrasonic Cell Disruptor, Emerson, Missouri, USA). The protein concentration of lysates was determined using BCA. After equalization, 50 mM Imidazole and 5 mM β-mercaptoethanol were added to each sample. Ni-NTA beads (QIAGEN) were used to enrich His10-Ubiquitin conjugates. Equilibrated Ni-NTA beads were added to the cell lysates and incubated o/n at 4°C under rotation.

Beads were pelleted at 1,000 g for 3 min and washed with the same buffers as described before in **Purification of TULIP2 conjugates**. The beads were transferred to an Eppendorf LoBind tube (Eppendorf) and washed for 15 min on a rotary wheel at RT with wash buffer 1, 2, 3 and 4. Ni-NTA beads were resuspended in 60 µL of Wash Buffer 5 and 10% of the sample was taken as pull-down for immunoblotting. The rest of the sample was prepared for mass spectrometry.

### iPOND

Isolation of Proteins On Nascent DNA (iPOND) technique was performed as established(*56*). Ten confluent 150 mm dishes of Parental and *RNF25^-/-^*cells were labeled with EdU for 15 min. Cells were fixed with 1% formaldehyde in PBS and cross-linking reaction was stopped with 1.25 M glycine for 5 min. Cells were pelleted, washed with cold PBS and incubated in permeabilization buffer (0.25% Triton X-100 in PBS) for 30 min. Cells were then washed with 0.5% BSA in PBS and incubated with a click reaction cocktail (PBS, 10 µM biotin azide (Invitrogen), 10 mM sodium ascorbate, 2 mM CuSO_4_ per 1 × 10^8^ cells) while rotating for 1–2 h. After the incubation time, cells were washed with cold 0.5% BSA and resuspended in lysis buffer [1% SDS in 50 mM Tris-HCl, (pH 8.0)] supplemented with aprotinin (Sigma-Aldrich) and leupeptin (Sigma-Aldrich). Lysates were sonicated (30 s constant pulse, 40 s pause at high power; total pulse time: 3 min) with a Bioruptor® Plus sonication device (Diagenode). Samples were centrifuged for 10 min at 16,000 g and the resulting lysate was diluted 1:1 (v/v) with cold PBS containing proteases inhibitors. Lysates were incubated with 100 µL of streptavidin beads (Sigma-Aldrich) slurry per 1 × 10^8^ cells for 16–20 h at 4°C while rotating. Beads were washed with cold lysis buffer and then transferred to an Eppendorf LoBind tube (Eppendorf) with Wash buffer A [0.1% sodium deoxycholate, 1% Triton X-100, 500 mM NaCl, 1 mM EDTA, 50 mM HEPES (pH 7.5)]. Subsequently, beads were washed with Wash buffer B [250 mM LiCl, 0.5% Triton X-100, 0.5% sodium deoxycholate, 1 mM EDTA, 10 mM Tris-HCl (pH 8)] and finally with Wash Buffer C [50 mM Tris-HCl (pH 7.5), 50 mM NaCl]. Samples were then prepared for mass spectrometry by washing the beads three times with 50 mM ammonium bicarbonate.

### TurboID

TurboID was performed as stablished^3^. 60 million cells per replicate of Parental constitutively expressing RNF25-WT-TurboID and C159/161S-TurboID, or Parental, *RNF25^-/-^*, and RNF25-GFP rescued cells inducibly expressing POLR2I-TurboID^3^ were used. POLR2I-TurboID construct was induced for 36 hours with 1 µg/mL or 200 ng/mL of doxycycline. Subsequently, cells were treated with 50 µM BioReagent-grade biotin (Sigma-Aldrich) for 10 min at 37°C, then washed with PBS, and crosslinked with 1% formaldehyde (Sigma-Aldrich) for 20 min at RT. Crosslinking reaction was quenched by adding 1.25 M glycine for 5 min and cells were scrapped and pelleted at 900 g for 5 min, washed with PBS and snap frozen. Pellets were resuspended in 2 mL of lysis buffer [50 mM Hepes (pH 7.9), 140 mM NaCl, 1 mM EDTA (pH 8), 10% glycerol, 0.5% IGEPAL, and 0.25% Triton X-100] containing protease inhibitors [cOmplete Mini EDTA-free Protease Inhibitor Cocktail (Roche)] and incubated for 10 min on ice. Cells nuclei were collected by centrifugating at 1,700 g for 5 min at 4°C and washed with [10 mM Tris-HCl (pH 8.0), 1 mM EDTA (pH 8.0), 0.5 mM EGTA (pH 8), 200 mM NaCl, and protease inhibitors] for 10 min on ice. After centrifugation, tubes were rinsed with shearing buffer [10 mM Tris-HCl (pH 7.6), 1 mM EDTA (pH 8), 0.1% SDS, and protease inhibitors], and the nuclei pellet was resuspended in shearing buffer and transferred to Adaptive Focused Acoustics (AFA) fiber 1-mL milliTUBE (Covaris). Chromatin was sheared using a Covaris ultrasonicator [Peak Incident Power (PIP): 140; duty: 5%; time: 900 s; 6°C]. After sonication, samples were centrifuged at 14,000 g for 10 min at 4°C, and the supernatant was transferred to a new tube for the streptavidin pull-down. 25 μL of Streptavidin beads (Millipore, 69203-3) were added per sample and incubated o/n at 4°C while rotating. After incubation, beads were washed with shearing buffer and decrosslinked with 1% SDS and 500 mM NaCl in TE buffer [10 mM Tris-HCl (pH 8) and 1 mM EDTA] at 65°C o/n at 600 rpm.

Samples were centrifuged at 500 g, and the supernatant was discarded. Beads were first washed with 2% SDS and then with same wash buffers detailed in **iPOND** methodology: Wash buffer A, B and C. Subsequently, samples were prepared for mass spectrometry by washing the beads three times with 50 mM ammonium bicarbonate.

### Digestion of purified conjugates

For protein digestion, samples were resuspended in 250 µL of ammonium bicarbonate (ABC) to be later subjected to reduction with 1 mM DTT for 30 mins, alkylation with 5 mM chloroacetamide for 20 mins and a further incubation with 5 mM DTT for 30 mins. Protein conjugates from Ni-NTA were first digested with 500 ng recombinant Lys-C (Promega) at RT while shaking at 1,400 rpm and subsequently diluted with 4 volumes of ABC. After digestion with Lys-C or not, protein conjugates were purified from Ni-NTA, GFP or streptavidin beads with 500 ng of sequencing grade modified trypsin (Promega) o/n at 37°C while shaking at 1,400 rpm. Trypsin digested peptides were separated from the beads by filtering through a 0.45 µm filter Ultrafree-MC-HV spin column (Merck-Millipore).

### Mass Spectrometry sample preparation

Digested peptides were acidified by adding 2% TriFlourAcetic (TFA) acid. Subsequently, peptides were desalted and concentrated on triple-disc C18 Stage-tips as previously described(*57*). Stage-tips were in-house assembled using 200 µL micro pipet tips and C18 matrix (Sigma Aldrich). Stage-tips were activated by passing through 100 µL of methanol. Next, 100 µL of Buffer A (80% acetonitrile, 0.1% formic acid) and 100 µL of Buffer B (0.1% formic acid) were passed through the activated C18 tip. Next, 100 µL the acidified peptide sample (this step was repeated until passing through the whole sample volume), and two times 100 µL Buffer B were passed through the Stage-tip. Elution was performed twice with 30 µL of Elution buffer (33% acetonitrile, 0.1% formic acid solution). Samples were vacuum dried using a Universal Vacuum System UVS400S coupled to a SpeedVac SPD121P (Thermo) and stored at -20°C. Prior to mass spectrometry analysis, samples were reconstituted in 10 or 20 µL 0.1% formic acid and transferred to autoload vials.

### LC-MS/MS data acquisition

The different mass spectrometry-based proteomics datasets were acquired using different equipment and acquisition settings depending on availability at the moment of performing the experiments. RNF25-TULIP2 data (Supplementary dataset 2, 3), RNF25 GFP-Trap CoIP (Supplementary dataset 4) and RNF25-TurboID (Supplementary dataset 5) and GFP-PCNA (Supplementary datasets 6,7) were acquired at the Proteomics facility in Institute of Biomedicine of Seville (IBiS). RNF25-TULIP2 and RNF25 GFP-Trap CoIP data were acquired in an Easy-nLC1000 liquid chromatrographer coupled to a Q-Exactive Plus mass spectrometer (Thermo Scientific) with a 100 minutes long chromatography gradient for the TULIP2 samples and 60 minutes for the GFP-Trap samples on a 500 mm Nanospray-Easy Chromatography column (Themo Scientific). In both cases from 2% to 33% Acetonitrile in 0.1% formic acid in water, followed to column re-equilibration and washing. For the mass spectrometer acquisition settings, a Top 15 Data Dependent Acquisition (DDA) method was used with a scan range of 400-1600 m/z and a resolution of 70,000 in the MS1. For the MS2, a resolution of 35,000, an AGC target of 5e5 and a Maximum Injection Time of 120 ms. Isolation window was 2.2 m/z and HCD fragmentation energy was set to 25%. A dynamic exclusion window of 60 s was applied. The RNF25-TurboID samples were measured using a nanoElute II LC system coupled to a timsTOF SCP mass spectrometer with an electrospray source (Bruker Daltonics). LC separations were performed on C18 HPLC column (Aurora 25 cm and 75 µm ID, IonOpticks) kept at 50°C. Gradient elution was performed with a binary system consisting of (A) 0.1% aqueous formic acid and (B) 0.1% formic acid in Acetonitrile. An increasing linear gradient (v/v) was used (t (min), %B): (0, 2); (40, 17); (60, 25); (66, 37); (67, 95), followed by an equilibration step. Mass spectrometric analysis was performed in a data independent acquisition parallel accumulation serial fragmentation (dia-PASEF) mode, with 100–1700 m/z mass range, an ion mobility range from 0.64 to 1.45 V s cm−2, capillary voltage set to 1500 V, an accumulation and ramp time at 100 ms and the collision energy as a linear ramp from 20 eV at 1/K0 = 0.6 V s cm−2 to 59 eV at 1/K0 = 1.6 V s cm^−2^.

The remaining datasets (Supplementary Datasets 1, 8, 9) were acquired at the proteomics service of CABIMER using a Vanquish Neo liquid chromatographer coupled to an Orbitrap Astral mass spectrometer (Thermo Scientific). Chromatography gradients separation was performed on a 15 cm PepMap Neo C18 analytical column on a Nanospray Flex Ion Source from 4% to 40% buffer B (80% Acetonitrile, 0.1% formic acid in water) in 23 minutes followed by column washing and equilibration. Spray Voltage was set at 2 kV and the Ion transfer tube temperature was 290°C. EASY-IC internal calibration was applied at RunStart. Acquisition was performed using Data Independent Acquisition (DIA) mode. The Master Scan was performed in the Orbitrap at a reolution of 240,000 and a normalized AGC target of 500%. DIA scans were performed in the Astral with a mass range from 380-980 m/z. For the DIA scans an isolation window of 4 Th and maximum injection time of 7 ms was used. The Normalized HCD Collision energy was set to 25% and Loop time was 0.6s

### Mass Spectrometry data analysis

RNF25-TULIP2 (Supplementary Dataset 2) raw data was analyzed using MaxQuant (version 2.6.7.0) as previously described (*58*). Search was performed against an in-silico digested UniProt reference proteome for *Homo sapiens* including canonical and isoform sequences (29^th^ August 2022). Database searches were performed according to standard settings with the following modifications: Digestion with Trypsin/P was used, allowing 3 missed cleavages. Oxidation (M) and GlyGly (K) for ubiquitination sites were allowed as variable modifications with a maximum number of 3. Label-Free Quantification (LFQ) was enabled, not allowing Fast LFQ while permitting iBAQ and matching between runs.

RNF25 GFP-Trap CoIP (Supplementary Dataset 4) raw data was analyzed using FragPipe v24 with MSFragger v4.4.1 (*59, 60*), IonQuant v1.11.20 (*61*) and DIA-NN v2.3.2 (*62*). An LFQ-MBR workflow was set, and samples data type was set to DDA+ (*63*). Reference proteome for Homo sapiens including canonical and isoform sequences (Uniprot 3^rd^ March 2026) was used. RNF25-TurboID (Supplementary Dataset 5) raw dia-PASEF data was analyzed using Biognosys Spectronaut (v 20.1.250624) using a directDIA+ search with BGS Factory settings. Predicted spectral library was built using a FASTA file corresponding to the reference human proteome (Uniprot 3^rd^ March 2026). A Pivot report was generated. Raw data from the rest of datasets (Supplementary Datasets 1, 6, 7, 8, 9) were analyzed using DIA-NN (v2.3.2 Academia) (*62*). A spectral library was *in silico* generated using the same FASTA file from the other datasets (Uniprot 3^rd^ March 2026) and the acquisition settings from the Orbitrap Astral mass spectrometer, namely, m/z ranges of 380-980 and 150-2000 for precursors and fragments, respectively. Charges between 2 and 5 and enabling a maximum of 2 missed cleavages for Trypsin.

The outputs from the different search engines were processed for statistical analysis in the Perseus computational platform (v.1.6.7.0)(*64*). LFQ intensity values were log2 transformed and, if applicable, potential contaminants and proteins were either identified by site or only reverse peptides were removed. Samples were grouped in experimental categories and proteins not identified in every replicate in at least one group were removed. Missing values were imputed using normally distributed values with 0.3 width and 1.8 down shift separately for each column. After imputation, statistical analysis was performed using two-sided Student’s t tests. Results were exported into MS Excel 365 for comprehensive browsing and visualization of the datasets. Volcano plots were constructed for data visualization using the VolcaNoseR web app(*65*) (https://huygens.science.uva.nl/VolcaNoseR/).

### Synthetic lethality CRISPR screening

Parental and *RNF25^-/-^* cells were transduced with the lentiviral Brunello-Cas9 library (Addgene, #73179-LV)(*49*) at a 300-fold coverage and ∼0.6 MOI. Puromycin-containing medium was added the next day to select for transductants. Three days after infection, which was considered the initial time point (T_0_), cells were pooled together and divided into two technical replicates, A and B. Negative-selection screens were performed by subculturing cells at day 3 (T_3_) and then subcultured every 3 days (T_6_, T_9_ and T_12_) until final time point at T_21_. Cell pellets were frozen at each time point for gDNA isolation. gDNA from cell pellets was isolated using the Wizard® Genomic DNA Purification Kit (Promega, A1120) and genome-integrated sgRNA sequences were amplified by PCR using Q5 Mastermix Next Ultra II (New England Biolabs, M5044L). Samples preparations were done amplifying the sgRNA with primers forward i5 and reverse i7 (see Supplementary Table 1 for primer list) and PCR products were purified using the Gel and PCR clean up Kit (Macherey-Nagel, 740609.250). Libraries were size-selected using dual selection with Sera-Mag beads at 0.7x-0.9x and were subsequently sequenced on Illumina NovaSeq 6000 system to determine sgRNA representation in each sample.

Demultiplexed reads were subsequently processed using MAGeCK v0.5.9.5 (*66*). Briefly, count subcommand was run to trim the 5’ end of the reads and to generate a raw counts file. Then, gene selection was assessed using the MLE algorithm, considering control sgRNAs for normalization between samples. Both Parental and RNF25-KO samples at 21 days were compared with their respective 0 days controls. Differential beta score was computed as RNF25 Knockout – Parental beta scores and sorted to obtain gene ranks. Genes displaying a differential beta score > 0.5 or < -0.5 were respectively classified as positive or negative selected. Visualization was conducted using custom R scripts under R v4.5.3.

### Clonogenic survival assay

Cells lines were seeded at 1,000 cells/well in 6-well plates. CHX, Olaparib, Methyl methanesulfonate or WEE1 inhibitor (MedChemExpress, HY-10993) were added the next day at different concentrations. After treatment, cells were allowed to grow for 7 days and fixed for 20 min in 4% paraformaldehyde (PFA) in PBS. Cells were stained with Crystal Violet 0.05% for 30 min and washed with water. Plates were scanned and colony formation was measured with Fiji plugin ColonyArea (*67*). GraphPad Prism 10 was used for statistical analysis, and the value of untreated cells was set at 100% survival.

### Immunofluorescence

Cell lines were seeded on 18 mm coverslips in 12-well plates. The next day, cells were treated with DMSO, 3 µg/mL CHX, 1 mM MMS for 3 hours, and irradiated with ultraviolet (UV) at 40 j/m^2^ dose with a recovery time of 5 hours. After treatment time, cells were fixed with 1% PFA, 0.3% Triton X-100 for 20 min and pre-extracted with 1% PFA, 0.3% Triton X-100 and 0.5% methanol for 20 min on a shaker at RT. Cells were blocked with TNB Blocking Reagent buffer (Roche) for 30 min and stained with primary antibodies diluted in TNB for 1.5 h at RT. Coverslips were washed four times with PBS and incubated with secondary antibodies in TNB for 1 h at RT. Next, coverslips were washed five times with PBS and mounted on slides with ProLong^TM^ Diamond Antifade Mountant with DAPI (Invitrogen, P36971). Imaging was performed using a confocal microscope (Leica SP5 Lightning) taking 8 frames per image. Around 50 cells per replicate were quantified using ImageJ Fiji (https://fiji.sc)(68) and data was represented using the SuperPlotsOfData website (https://huygens.science.uva.nl/SuperPlotsOfData)(69). Statistical details of individual immunofluorescence experiments can be found in figure legends. Antibodies are listed in Supplementary Tables 3 and 4.

### S1 nuclease DNA fiber assay

S1 nuclease DNA fiber assay was followed as described^2^. Cells were pulse-labeled with 20 µM IdU (20 min), washed and pulse-labeled with 200 µM CldU and treated or not with 3 µg/mL CHX for 1 h. Cells were then washed twice and permeabilized with CSK buffer [100 mM NaCl, 10 mM MOPS (pH 7), 3 mM MgCl_2_, 300 mM Sucrose, and 0.5% Triton X-100 in water] for 8 min at RT. Permeabilized cells were treated with S1 nuclease buffer [30 mM Sodium acetate (pH 4.6), 10 mM Zinc acetate, 5% glycerol, 50 mM NaCl in water] with or without 20 U/mL S1 nuclease (Invitrogen, 18001-016) for 30 min at 37°C. Subsequently, cells were scrapped in PBS 0.1% BSA, pelleted, and resuspended in PBS 0.1% BSA at a final concentration of 1-2 × 10^3^ cells/µl. Cell suspension of 2.5 µL was spotted on a positively charged slide (Epredia, J1800AMNZ) and lysed with 7.5 µL of spreading buffer [200 mM Tris-HCl (pH 7.5), 50 mM EDTA, 0.5% SDS]. After 8 min, slides were tilted at 45 degrees to allow the DNA to spread. Slides were air-dried, fixed with ice-cold methanol/acetic acid (3:1) for 5 mins, air-dried, and stored at 4°C until further staining. Slides were rehydrated with PBS, denatured with 2.5 M HCl for 1 h, washed twice, and blocked with blocking buffer (3% BSA, 0.1% Triton X-100 in PBS) for 30 min. Next, slides were incubated with primary antibodies diluted in blocking buffer for 2.5 h at RT in a dark humid chamber and washed three times with PBS. Slides were then incubated with secondary antibodies in blocking buffer for 1 h at RT in a dark humid chamber. After washing and air-drying, slides were mounted with ProLong^TM^ Gold antifade reagent (Invitrogen, P36930). Images were acquired using AF6000 Leica Fluorescence microscope equipped with a HCX PL APO 63× (NA = 1.4) oil objective. At least 100 fibers per condition were measured using the segmented line tool on ImageJ Fiji (https://fiji.sc)(68). Statistical details of S1 DNA fiber experiments can be found in figure legends; these were performed according to the methods protocol published by Vindigni lab(*70*). Antibodies are listed in Supplementary Tables 3 and 4.

### Omics data availability

The mass spectrometry proteomics data have been deposited to the ProteomeXchange Consortium via the PRIDE (*71*) partner repository with the dataset identifier PXD078282.

For reviewing purposes, the following credentials can be used:

**Project accession:** PXD078282

**Token:** vPlG1Sq377wV

The DNA sequencing data from the CRISPR/Cas9 screens have been deposited in the Gene Expression Omnibus (*72*) with the accession number GSE332950.

**Revier token:** ctmrewgkrvenbyl

## ACKNOWLEDGEMENTS

Authors thanks Sylvie Noordeemer for providing the TP53^-/-^ PAC^-/-^ RPE1 (*73*) cells used in this study as Parental Cells and Andrés Aguilera and Emilia Herrera-Moyano for sharing the inducible RNAaseH1-eGFP plasmid.

Research and publication of this work was supported by the project PID2024-159761NB-I00 financed by MICIU/AEI/10.13039/501100011033 and FEDER, UE to RG-P.

Additional sources of funding for research include: EMERGIA 2020 program (EMERGIA20_00276) from the Consejería de Economía, Conocimiento, Empresas y Universidad, Junta de Andalucía, Spain and grants CNS2022-135216 funded by MICIU/AEI/10.13039/501100011033 and by European Union NextGenerationEU/PRTR and PID2021-122361NA-I00 by MICIU/AEI/10.13039/501100011033 and by European Union to RG-P. The N.G-R. laboratory was supported by grant PID2023-150072N, funded by MICIU/AEI/10.13039/501100011033 and by ERDF/EU; by grant CNS2024-154523 funded by MICIU/AEI/10.13039/501100011033; and by the EMERGIA 2021 program from the Junta de Andalucía (EMC21_00057). Grant PID2022-139691OB-I00 funded by MICIU/AEI/10.13039/501100011033 and ERDF/EU to RF. The GM-Z laboratory was supported by grants PID2021-127432NA-I00 and PID2023-151942NB-I00 funded by MICIU/AEI/10.13039/501100011033 and by ERDF/EU, by grant CNS2022-135600 funded by MICIU/AEI/10.13039/501100011033 and by European Union NextGenerationEU/PRTR

CE-S is supported by Juan de la Cierva program, grant JDC2023-051129-I financed by MICIU/AEI/10.13039/501100011033 and FSE+. LG-V is supported by the Spanish Ministry of Science, Innovation and Universities, fellowship FPU2023/03611.

## AUTHOR CONTRIBUTION

ES-H, CE-S, LG-V and MM-C performed experiments. ES-H, RdA and RG-P analyzed experimental data. MLM-M and ARH acquired mass spectrometry data. NG-R trained ES-H in DNA fiber assays. RF supervised MM-C. GM-Z supervised RdA. RG-P supervised ES-H, CE-S, LG-V and ARH. ES-H and RG-P wrote the manuscript with input the rest of authors.

## REFERENCES

1. D. Salas-Lloret, R. Gonzalez-Prieto, Insights in Post-Translational Modifications: Ubiquitin and SUMO. Int J Mol Sci 23, (2022).

2. M. Olivieri, T. Cho, A. Alvarez-Quilon, K. Li, M. J. Schellenberg, M. Zimmermann, N. Hustedt, S. E. Rossi, S. Adam, H. Melo, A. M. Heijink, G. Sastre-Moreno, N. Moatti, R. K. Szilard, A. McEwan, A. K. Ling, A. Serrano-Benitez, T. Ubhi, S. Feng, J. Pawling, I. Delgado-Sainz, M. W. Ferguson, J. W. Dennis, G. W. Brown, F. Cortes-Ledesma, R. S. Williams, A. Martin, D. Xu, D. Durocher, A Genetic Map of the Response to DNA Damage in Human Cells. Cell 182, 481–496 e421 (2020).

3. S. M. Noordermeer, S. Adam, D. Setiaputra, M. Barazas, S. J. Pettitt, A. K. Ling, M. Olivieri, A. Alvarez-Quilon, N. Moatti, M. Zimmermann, S. Annunziato, D. B. Krastev, F. Song, I. Brandsma, J. Frankum, R. Brough, A. Sherker, S. Landry, R. K. Szilard, M. M. Munro, A. McEwan, T. Goullet de Rugy, Z. Y. Lin, T. Hart, J. Moffat, A. C. Gingras, A. Martin, H. van Attikum, J. Jonkers, C. J. Lord, S. Rottenberg, D. Durocher, The shieldin complex mediates 53BP1-dependent DNA repair. Nature 560, 117–121 (2018).

4. N. Y. L. Ngoi, D. Gallo, C. Torrado, M. Nardo, D. Durocher, T. A. Yap, Synthetic lethal strategies for the development of cancer therapeutics. Nat Rev Clin Oncol 22, 46–64 (2025).

5. Y. van der Weegen, K. de Lint, D. van den Heuvel, Y. Nakazawa, T. E. T. Mevissen, J. J. M. van Schie, M. San Martin Alonso, D. E. C. Boer, R. Gonzalez-Prieto, I. V. Narayanan, N. H. M. Klaassen, A. P. Wondergem, K. Roohollahi, J. C. Dorsman, Y. Hara, A. C. O. Vertegaal, J. de Lange, J. C. Walter, S. M. Noordermeer, M. Ljungman, T. Ogi, R. M. F. Wolthuis, M. S. Luijsterburg, ELOF1 is a transcription-coupled DNA repair factor that directs RNA polymerase II ubiquitylation. Nature cell biology 23, 595–607 (2021).

6. D. van den Heuvel, M. Rodriguez-Martinez, P. J. van der Meer, N. Nieto Moreno, J. Park, H. S. Kim, J. J. M. van Schie, A. P. Wondergem, A. D’Souza, G. Yakoub, A. E. Herlihy, K. Kashyap, T. Boissiere, J. Walker, R. Mitter, K. Apelt, K. de Lint, I. Kirdok, M. Ljungman, R. M. F. Wolthuis, P. Cramer, O. D. Scharer, G. Kokic, J. Q. Svejstrup, M. S. Luijsterburg, STK19 facilitates the clearance of lesion-stalled RNAPII during transcription-coupled DNA repair. Cell 187, 7107–7125 e7125 (2024).

7. S. Zhao, C. S. Palma-Chaundler, C. M. Engel, J. Cordes, D. Nixdorf, M. Y. Luo, S. Kaya, A. Suryo Rahmanto, D. van den Heuvel, T. Mackens-Kiani, P. Weickert, S. Lam, V. Gupta, J. Philippou-Massier, I. Bagaric, J. Bohlen, G. Hewitt, M. S. Luijsterburg, R. Beckmann, P. Beli, D. D. Nedialkova, C. J. Carnie, M. Subklewe, S. P. Jackson, J. Stingele, RNF25 confers mRNA damage tolerance by curbing activation of the integrated stress response. Molecular cell, (2026).

8. L. F. Chiou, D. Jayaprakash, G. N. Droby, X. Zhang, Y. Yang, C. A. Mills, T. S. Webb, N. K. Barker, D. Wu, L. E. Herring, J. Bowser, C. Vaziri, The RING Finger E3 Ligase RNF25 Protects DNA Replication Forks Independently of its Canonical Roles in Ubiquitin Signaling. bioRxiv, (2025).

9. L. F. Chiou, G. N. Droby, D. Jayaprakash, J. R. Anand, X. Zhang, Y. Yang, C. A. Mills, T. S. Webb, N. K. Barker, J. Xie, D. Wu, L. E. Herring, J. Tomida, J. L. Bowser, C. Vaziri, The RING finger E3 ligase RNF25 protects DNA replication forks independently of its canonical roles in ubiquitin signaling. Nature communications 16, 7214 (2025).

10. S. Zhao, J. Cordes, K. M. Caban, M. J. Gotz, T. Mackens-Kiani, A. J. Veltri, N. K. Sinha, P. Weickert, S. Kaya, G. Hewitt, D. D. Nedialkova, T. Frohlich, R. Beckmann, A. R. Buskirk, R. Green, J. Stingele, RNF14-dependent atypical ubiquitylation promotes translation-coupled resolution of RNA-protein crosslinks. Molecular cell 83, 4290–4303 e4299 (2023).

11. K. Oltion, J. D. Carelli, T. Yang, S. K. See, H. Y. Wang, M. Kampmann, J. Taunton, An E3 ligase network engages GCN1 to promote the degradation of translation factors on stalled ribosomes. Cell 186, 346–362 e317 (2023).

12. C. Zhang, C. Tian, R. Zhu, C. Chen, C. Jin, X. Wang, L. Sun, W. Peng, D. Ji, Y. Zhang, Y. Sun, CircSATB1 Promotes Colorectal Cancer Liver Metastasis through Facilitating FKBP8 Degradation via RNF25-Mediated Ubiquitination. Adv Sci (Weinh) 12, e2406962 (2025).

13. L. Li, Z. Wang, B. Ma, Q. Ye, Y. Lei, M. Lu, L. Ye, J. Kang, W. Huang, S. Xu, K. Wang, Y. Chen, J. Liu, Y. Gao, C. Wang, J. Ma, L. Li, BAY11-7082 Targets RNF25 to Reverse TRIP4 Ubiquitination-dependent NF-kappaB Activation and Apoptosis Resistance in Renal Cell Carcinoma. International journal of biological sciences 21, 4410–4427 (2025).

14. Z. Huang, L. Zhou, J. Duan, S. Qin, J. Jiang, H. Chen, K. Wang, R. Liu, M. Yuan, X. Tang, E. C. Nice, Y. Wei, W. Zhang, C. Huang, Oxidative Stress Promotes Liver Cancer Metastasis via RNF25-Mediated E-Cadherin Protein Degradation. Adv Sci (Weinh) 11, e2306929 (2024).

15. J. H. Cho, Y. M. You, Y. I. Yeom, D. C. Lee, B. K. Kim, M. Won, B. C. Cho, M. Kang, S. Park, S. J. Yang, J. S. Kim, J. A. Kim, K. C. Park, RNF25 promotes gefitinib resistance in EGFR-mutant NSCLC cells by inducing NF-kappaB-mediated ERK reactivation. Cell Death Dis 9, 587 (2018).

16. N. Tsao, J. R. Brickner, R. Rodell, A. Ganguly, M. Wood, C. Oyeniran, T. Ahmad, H. Sun, A. Bacolla, L. Zhang, V. Lukinovic, J. M. Soll, B. A. Townley, A. G. Casanova, J. A. Tainer, C. He, A. Vindigni, N. Reynoird, N. Mosammaparast, Aberrant RNA methylation triggers recruitment of an alkylation repair complex. Molecular cell 81, 4228–4242 e4228 (2021).

17. J. Cordes, S. Zhao, C. M. Engel, J. Stingele, Cellular responses to RNA damage. Cell 188, 885–900 (2025).

18. A. C. Vind, Z. Wu, M. J. Firdaus, G. Snieckute, G. A. Toh, M. Jessen, J. F. Martinez, P. Haahr, T. L. Andersen, M. Blasius, L. F. Koh, N. L. Maartensson, J. E. A. Common, M. Gyrd-Hansen, F. L. Zhong, S. Bekker-Jensen, The ribotoxic stress response drives acute inflammation, cell death, and epidermal thickening in UV-irradiated skin in vivo. Molecular cell 84, 4774–4789 e4779 (2024).

19. N. K. Sinha, C. McKenney, Z. Y. Yeow, J. J. Li, K. H. Nam, T. M. Yaron-Barir, J. L. Johnson, E. M. Huntsman, L. C. Cantley, A. Ordureau, S. Regot, R. Green, The ribotoxic stress response drives UV-mediated cell death. Cell 187, 3652–3670 e3640 (2024).

20. L. Wan, S. Juszkiewicz, D. Blears, P. K. Bajpe, Z. Han, P. Faull, R. Mitter, A. Stewart, A. P. Snijders, R. S. Hegde, J. Q. Svejstrup, Translation stress and collided ribosomes are co-activators of cGAS. Molecular cell 81, 2808–2822 e2810 (2021).

21. D. Salas-Lloret, N. Garcia-Rodriguez, E. Soto-Hidalgo, L. Gonzalez-Vinceiro, C. Espejo-Serrano, L. Giebel, M. L. Mateos-Martin, A. H. de Ru, P. A. van Veelen, P. Huertas, A. C. O. Vertegaal, R. Gonzalez-Prieto, BRCA1/BARD1 ubiquitinates PCNA in unperturbed conditions to promote continuous DNA synthesis. Nature communications 15, 4292 (2024).

22. D. Salas-Lloret, G. Agabitini, R. Gonzalez-Prieto, TULIP2: An Improved Method for the Identification of Ubiquitin E3-Specific Targets. Front Chem 7, 802 (2019).

23. Z. Yalcin, D. Koot, K. Bezstarosti, D. Salas-Lloret, O. B. Bleijerveld, V. Boersma, M. Falcone, R. Gonzalez-Prieto, M. Altelaar, J. A. A. Demmers, J. J. L. Jacobs, Ubiquitinome Profiling Reveals in Vivo UBE2D3 Targets and Implicates UBE2D3 in Protein Quality Control. Molecular & cellular proteomics : MCP 22, 100548 (2023).

24. S. Li, Y. H. Liang, J. Mariano, M. B. Metzger, D. K. Stringer, V. A. Hristova, J. Li, P. A. Randazzo, Y. C. Tsai, X. Ji, A. M. Weissman, Insights into Ubiquitination from the Unique Clamp-like Binding of the RING E3 AO7 to the E2 UbcH5B. The Journal of biological chemistry 290, 30225–30239 (2015).

25. Y. David, T. Ziv, A. Admon, A. Navon, The E2 ubiquitin-conjugating enzymes direct polyubiquitination to preferred lysines. The Journal of biological chemistry 285, 8595–8604 (2010).

26. H. J. Meyer, M. Rape, Enhanced protein degradation by branched ubiquitin chains. Cell 157, 910–921 (2014).

27. N. Garcia-Rodriguez, R. P. Wong, H. D. Ulrich, Functions of Ubiquitin and SUMO in DNA Replication and Replication Stress. Frontiers in genetics 7, 87 (2016).

28. M. Galloy, A. Blondeau, E. Vion, D. Kim, C. A. Bakker, V. Gaggioli, M. Thomas, E. G. Lavoie, I. Marois, A. D. Delgado Monterroso, J. Y. Masson, N. Taneja, K. M. Miller, A. Marechal, A. Fradet-Turcotte, Ubiquitination of the histone variant mH2A1.2 prevents toxic RAD18 accumulation at a subset of genomic loci upon replication stress. Molecular cell, (2025).

29. T. Thakar, W. Leung, C. M. Nicolae, K. E. Clements, B. Shen, A. K. Bielinsky, G. L. Moldovan, Ubiquitinated-PCNA protects replication forks from DNA2-mediated degradation by regulating Okazaki fragment maturation and chromatin assembly. Nature communications 11, 2147 (2020).

30. S. Nayak, J. A. Calvo, K. Cong, M. Peng, E. Berthiaume, J. Jackson, A. M. Zaino, A. Vindigni, M. K. Hadden, S. B. Cantor, Inhibition of the translesion synthesis polymerase REV1 exploits replication gaps as a cancer vulnerability. Sci Adv 6, eaaz7808 (2020).

31. Z. Yu, S. Y. Mersaoui, L. Guitton-Sert, Y. Coulombe, J. Song, J. Y. Masson, S. Richard, DDX5 resolves R-loops at DNA double-strand breaks to promote DNA repair and avoid chromosomal deletions. NAR Cancer 2, zcaa028 (2020).

32. M. Polenkowski, A. B. Allister, S. Burbano de Lara, A. Pierce, B. Geary, O. El Bounkari, L. Wiehlmann, A. Hoffmann, A. D. Whetton, T. Tamura, D. D. H. Tran, THOC5 complexes with DDX5, DDX17, and CDK12 to regulate R loop structures and transcription elongation rate. iScience 26, 105784 (2023).

33. B. Boleslavska, A. Oravetzova, K. Shukla, Z. Nascakova, O. N. Ibini, Z. Hasanova, M. Andrs, R. Kanagaraj, J. Dobrovolna, P. Janscak, DDX17 helicase promotes resolution of R-loop-mediated transcription-replication conflicts in human cells. Nucleic acids research 50, 12274–12290 (2022).

34. M. Li, X. Xu, C. W. Chang, Y. Liu, TRIM28 functions as the SUMO E3 ligase for PCNA in prevention of transcription induced DNA breaks. Proceedings of the National Academy of Sciences of the United States of America 117, 23588–23596 (2020).

35. B. Singh, S. Roy Chowdhury, M. S. Mansuri, S. J. Pillai, S. Mehrotra, The BRCA2 and CDKN1A-interacting protein (BCCIP) stabilizes stalled replication forks and prevents degradation of nascent DNA. FEBS Lett 596, 2041–2055 (2022).

36. H. M. Ng, L. Wei, L. Lan, M. S. Huen, The Lys63-deubiquitylating Enzyme BRCC36 Limits DNA Break Processing and Repair. The Journal of biological chemistry 291, 16197–16207 (2016).

37. K. A. Coleman, R. A. Greenberg, The BRCA1-RAP80 complex regulates DNA repair mechanism utilization by restricting end resection. The Journal of biological chemistry 286, 13669–13680 (2011).

38. A. Kakarougkas, A. Ismail, Y. Katsuki, R. Freire, A. Shibata, P. A. Jeggo, Co-operation of BRCA1 and POH1 relieves the barriers posed by 53BP1 and RAP80 to resection. Nucleic acids research 41, 10298–10311 (2013).

39. J. Sun, K. J. Lee, A. J. Davis, D. J. Chen, Human Ku70/80 protein blocks exonuclease 1-mediated DNA resection in the presence of human Mre11 or Mre11/Rad50 protein complex. The Journal of biological chemistry 287, 4936–4945 (2012).

40. M. Berti, A. Ray Chaudhuri, S. Thangavel, S. Gomathinayagam, S. Kenig, M. Vujanovic, F. Odreman, T. Glatter, S. Graziano, R. Mendoza-Maldonado, F. Marino, B. Lucic, V. Biasin, M. Gstaiger, R. Aebersold, J. M. Sidorova, R. J. Monnat, Jr., M. Lopes, A. Vindigni, Human RECQ1 promotes restart of replication forks reversed by DNA topoisomerase I inhibition. Nature structural & molecular biology 20, 347–354 (2013).

41. H. Stoy, K. Zwicky, D. Kuster, K. S. Lang, J. Krietsch, M. P. Crossley, J. A. Schmid, K. A. Cimprich, H. Merrikh, M. Lopes, Direct visualization of transcription-replication conflicts reveals post-replicative DNA:RNA hybrids. Nature structural & molecular biology 30, 348–359 (2023).

42. L. Gonzalez-Vinceiro, C. Espejo-Serrano, M. E. Soler-Oliva, E. Soto-Hidalgo, M. L. Mateos-Martin, D. Rico, C. Gonzalez-Aguilera, R. Gonzalez-Prieto, PLAMseq enables the proteo-genomic characterization of chromatin-associated proteins and protein interactions in a single workflow. Sci Adv 11, eady4151 (2025).

43. D. M. Moquin, M. M. Genois, J. M. Zhang, J. Ouyang, T. Yadav, R. Buisson, S. A. Yazinski, J. Tan, M. Boukhali, J. P. Gagne, G. G. Poirier, L. Lan, W. Haas, L. Zou, Localized protein biotinylation at DNA damage sites identifies ZPET, a repressor of homologous recombination. Genes & development 33, 75–89 (2019).

44. K. J. Abraham, J. N. Chan, J. S. Salvi, B. Ho, A. Hall, E. Vidya, R. Guo, S. A. Killackey, N. Liu, J. E. Lee, G. W. Brown, K. Mekhail, Intersection of calorie restriction and magnesium in the suppression of genome-destabilizing RNA-DNA hybrids. Nucleic acids research 44, 8870–8884 (2016).

45. S. H. Hung, R. P. Wong, H. D. Ulrich, C. F. Kao, Monoubiquitylation of histone H2B contributes to the bypass of DNA damage during and after DNA replication. Proceedings of the National Academy of Sciences of the United States of America 114, E2205–E2214 (2017).

46. B. Benedict, M. A. van Bueren, F. P. van Gemert, C. Lieftink, S. Guerrero Llobet, M. A. van Vugt, R. L. Beijersbergen, H. Te Riele, The RECQL helicase prevents replication fork collapse during replication stress. Life Sci Alliance 3, (2020).

47. A. Taglialatela, S. Alvarez, G. Leuzzi, V. Sannino, L. Ranjha, J. W. Huang, C. Madubata, R. Anand, B. Levy, R. Rabadan, P. Cejka, V. Costanzo, A. Ciccia, Restoration of Replication Fork Stability in BRCA1- and BRCA2-Deficient Cells by Inactivation of SNF2-Family Fork Remodelers. Molecular cell 68, 414–430 e418 (2017).

48. K. R. Sanson, R. E. Hanna, M. Hegde, K. F. Donovan, C. Strand, M. E. Sullender, E. W. Vaimberg, A. Goodale, D. E. Root, F. Piccioni, J. G. Doench, Optimized libraries for CRISPR-Cas9 genetic screens with multiple modalities. Nature communications 9, 5416 (2018).

49. J. G. Doench, N. Fusi, M. Sullender, M. Hegde, E. W. Vaimberg, K. F. Donovan, I. Smith, Z. Tothova, C. Wilen, R. Orchard, H. W. Virgin, J. Listgarten, D. E. Root, Optimized sgRNA design to maximize activity and minimize off-target effects of CRISPR-Cas9. Nature biotechnology 34, 184–191 (2016).

50. N. Minsky, E. Shema, Y. Field, M. Schuster, E. Segal, M. Oren, Monoubiquitinated H2B is associated with the transcribed region of highly expressed genes in human cells. Nature cell biology 10, 483–488 (2008).

51. S. J. Smith, F. Meng, R. G. Lingeman, C. M. Li, M. Li, G. Boneh, T. T. Seppala, T. Phan, H. Li, R. A. Burkhart, V. Parekh, S. Rahmanuddin, L. G. Melstrom, R. J. Hickey, V. Chung, Y. Liu, L. H. Malkas, M. Raoof, Therapeutic Targeting of Oncogene-Induced Transcription-Replication Conflicts in Pancreatic Ductal Adenocarcinoma. Gastroenterology 169, 600–614 e611 (2025).

52. A. da Costa, D. Chowdhury, G. I. Shapiro, A. D. D’Andrea, P. A. Konstantinopoulos, Targeting replication stress in cancer therapy. Nat Rev Drug Discov 22, 38–58 (2023).

53. L. González-Vinceiro, C. Espejo-Serrano, M. E. Soler-Oliva, M. L. Mateos-Martín, D. Rico, C. González-Aguilera, R. González Prieto, PLAMseq enables the proteo-genomic characterization of chromatin-associated proteins and protein interactions in a single experimental workflow. bioRxiv, 2025.2004.2027.650851 (2025).

54. X. Yang, J. S. Boehm, X. Yang, K. Salehi-Ashtiani, T. Hao, Y. Shen, R. Lubonja, S. R. Thomas, O. Alkan, T. Bhimdi, T. M. Green, C. M. Johannessen, S. J. Silver, C. Nguyen, R. R. Murray, H. Hieronymus, D. Balcha, C. Fan, C. Lin, L. Ghamsari, M. Vidal, W. C. Hahn, D. E. Hill, D. E. Root, A public genome-scale lentiviral expression library of human ORFs. Nature methods 8, 659–661 (2011).

55. N. Krupka, P. Strappe, J. Gotz, L. M. Ittner, Gateway-compatible lentiviral transfer vectors for ubiquitin promoter driven expression of fluorescent fusion proteins. Plasmid 63, 155–160 (2010).

56. B. M. Sirbu, F. B. Couch, D. Cortez, Monitoring the spatiotemporal dynamics of proteins at replication forks and in assembled chromatin using isolation of proteins on nascent DNA. Nat Protoc 7, 594–605 (2012).

57. J. Rappsilber, M. Mann, Y. Ishihama, Protocol for micro-purification, enrichment, pre-fractionation and storage of peptides for proteomics using StageTips. Nat Protoc 2, 1896–1906 (2007).

58. S. Tyanova, T. Temu, J. Cox, The MaxQuant computational platform for mass spectrometry-based shotgun proteomics. Nat Protoc 11, 2301–2319 (2016).

59. A. T. Kong, F. V. Leprevost, D. M. Avtonomov, D. Mellacheruvu, A. I. Nesvizhskii, MSFragger: ultrafast and comprehensive peptide identification in mass spectrometry-based proteomics. Nature methods 14, 513–520 (2017).

60. G. C. Teo, D. A. Polasky, F. Yu, A. I. Nesvizhskii, Fast Deisotoping Algorithm and Its Implementation in the MSFragger Search Engine. Journal of proteome research 20, 498–505 (2021).

61. F. Yu, S. E. Haynes, A. I. Nesvizhskii, IonQuant Enables Accurate and Sensitive Label-Free Quantification With FDR-Controlled Match-Between-Runs. Molecular & cellular proteomics : MCP 20, 100077 (2021).

62. V. Demichev, C. B. Messner, S. I. Vernardis, K. S. Lilley, M. Ralser, DIA-NN: neural networks and interference correction enable deep proteome coverage in high throughput. Nature methods 17, 41–44 (2020).

63. F. Yu, Y. Deng, A. I. Nesvizhskii, MSFragger-DDA+ enhances peptide identification sensitivity with full isolation window search. Nature communications 16, 3329 (2025).

64. S. Tyanova, T. Temu, P. Sinitcyn, A. Carlson, M. Y. Hein, T. Geiger, M. Mann, J. Cox, The Perseus computational platform for comprehensive analysis of (prote)omics data. Nature methods 13, 731–740 (2016).

65. J. Goedhart, M. S. Luijsterburg, VolcaNoseR is a web app for creating, exploring, labeling and sharing volcano plots. Sci Rep 10, 20560 (2020).

66. W. Li, J. Koster, H. Xu, C. H. Chen, T. Xiao, J. S. Liu, M. Brown, X. S. Liu, Quality control, modeling, and visualization of CRISPR screens with MAGeCK-VISPR. Genome biology 16, 281 (2015).

67. C. Guzman, M. Bagga, A. Kaur, J. Westermarck, D. Abankwa, ColonyArea: an ImageJ plugin to automatically quantify colony formation in clonogenic assays. PLoS One 9, e92444 (2014).

68. J. Schindelin, I. Arganda-Carreras, E. Frise, V. Kaynig, M. Longair, T. Pietzsch, S. Preibisch, C. Rueden, S. Saalfeld, B. Schmid, J. Y. Tinevez, D. J. White, V. Hartenstein, K. Eliceiri, P. Tomancak, A. Cardona, Fiji: an open-source platform for biological-image analysis. Nature methods 9, 676-682 (2012).

69. J. Goedhart, SuperPlotsOfData-a web app for the transparent display and quantitative comparison of continuous data from different conditions. Molecular biology of the cell 32, 470–474 (2021).

70. A. Quinet, D. Carvajal-Maldonado, D. Lemacon, A. Vindigni, DNA Fiber Analysis: Mind the Gap! Methods in enzymology 591, 55–82 (2017).

71. Y. Perez-Riverol, C. Bandla, D. J. Kundu, S. Kamatchinathan, J. Bai, S. Hewapathirana, N. S. John, A. Prakash, M. Walzer, S. Wang, J. A. Vizcaino, The PRIDE database at 20 years: 2025 update. Nucleic acids research 53, D543–D553 (2025).

72. T. Barrett, S. E. Wilhite, P. Ledoux, C. Evangelista, I. F. Kim, M. Tomashevsky, K. A. Marshall, K. H. Phillippy, P. M. Sherman, M. Holko, A. Yefanov, H. Lee, N. Zhang, C. L. Robertson, N. Serova, S. Davis, A. Soboleva, NCBI GEO: archive for functional genomics data sets--update. Nucleic acids research 41, D991–995 (2013).

73. B. van de Kooij, A. Schreuder, R. S. Pavani, V. Garzero, A. Van Hoeck, M. San Martin Alonso, D. Koerse, T. J. Wendel, E. Callen, J. Boom, H. Mei, E. Cuppen, A. Nussenzweig, H. van Attikum, S. M. Noordermeer, EXO1-mediated DNA repair by single-strand annealing is essential for BRCA1-deficient cells. bioRxiv, 2023.2002.2024.529205 (2023).

