## Supplemental Data for "RNF25 Ubiquitin E3 activity Safeguards Genome Integrity by Modulating DNA Replication, Transcription and Translation"

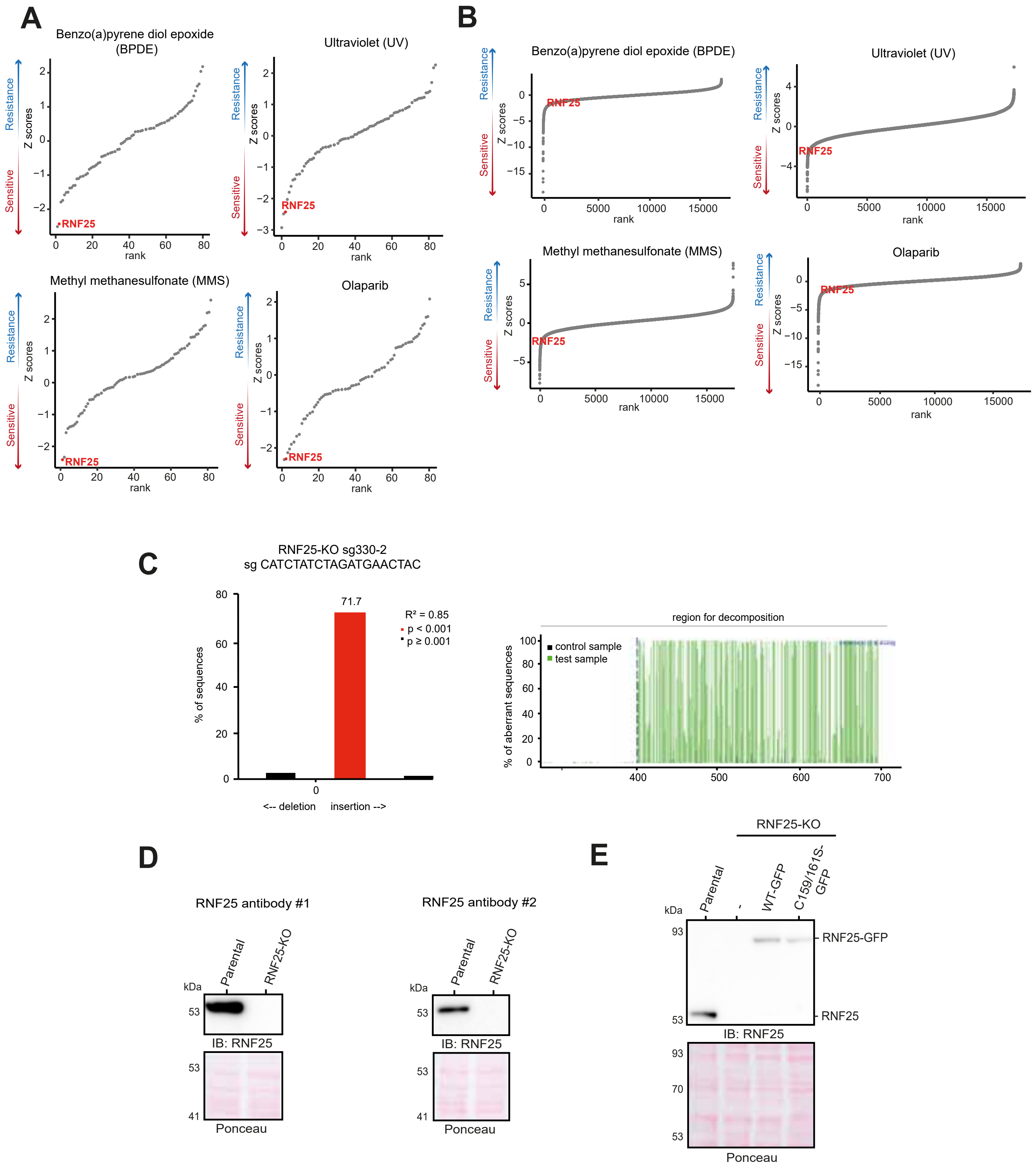


**Supplementary Figure 1. (A, B)** Gene ranks with Z scores from Daniel Durocher's laboratory genome wide CRISPR/Cas9 screening results, under treatment with Benzo(a)pyrene diol epoxide (BPDE), Ultraviolet radiation (UV), Methyl methanesulfonate (MMS) and Olaparib, focusing on RING Finger ubiquitin ligases **(A)** or all genes **(B)**. Each dot represents a gene. RNF25 is highlighted in red. **(C)** RNF25 knockout (KO) mutation characterization with TIDE tool. **(D)** Immunoblotting of Parental and RNF25-KO cells with two anti-RNF25 antibodies. **(E)** Immunoblotting of Parental and RNF25-KO cells rescued or not with RNF25-WT-GFP and catalytic dead mutant RNF25-C159/161S-GFP. Ponceau staining is shown as loading control.

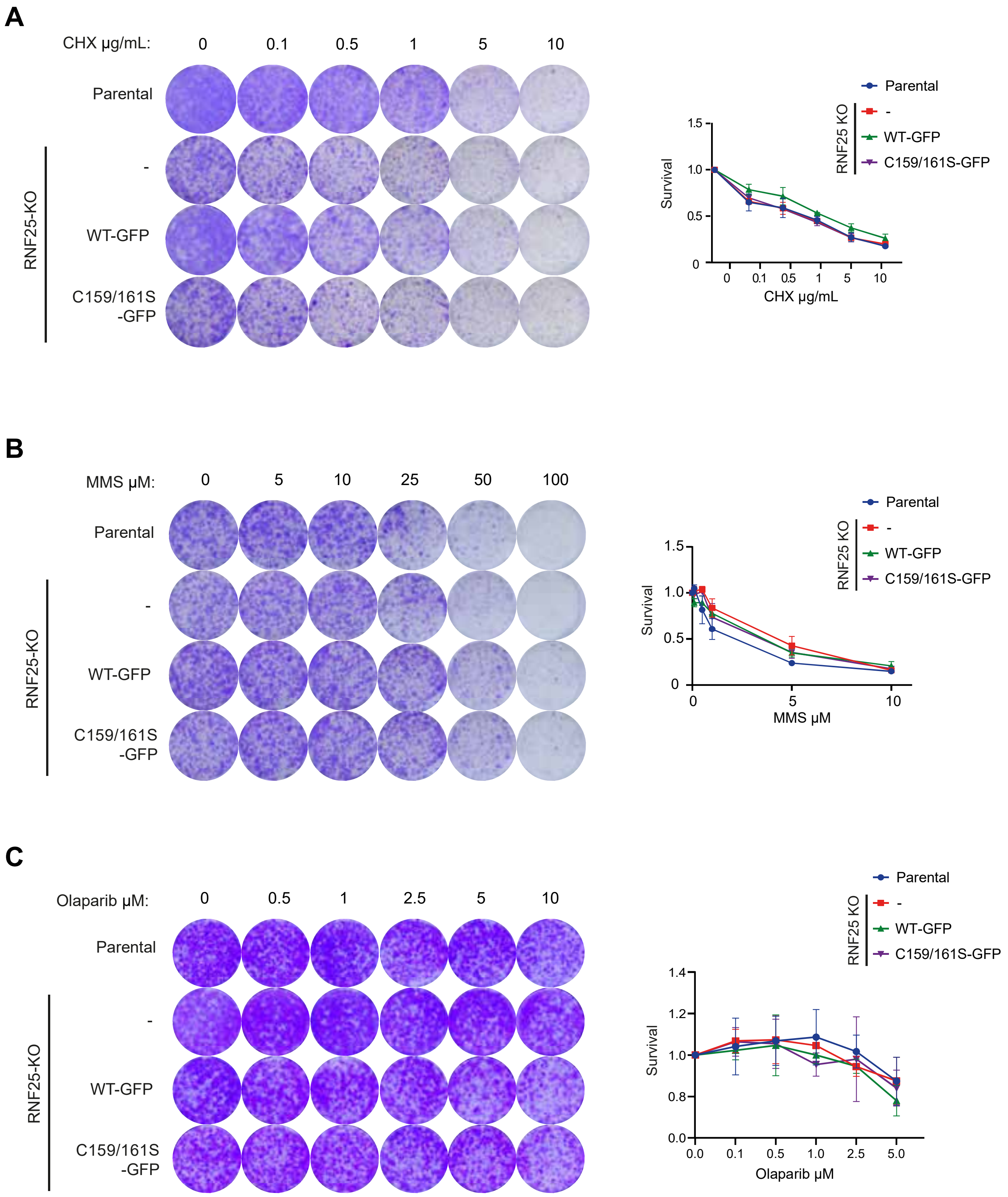


**Supplementary Figure 2. (A-C)** Clonogenic survival assay of Parental and RNF25-KO cells rescued or not with RNF25-WT-GFP or RNF25-C159/161S-GFP, under treatment with Methyl methanesulfonate (MMS), Cycloheximide (CHX) and Olaparib for 24 hours and 7-days recovery. Average and standard deviation of three independent experiments with three technical repeats are shown (N = 3).

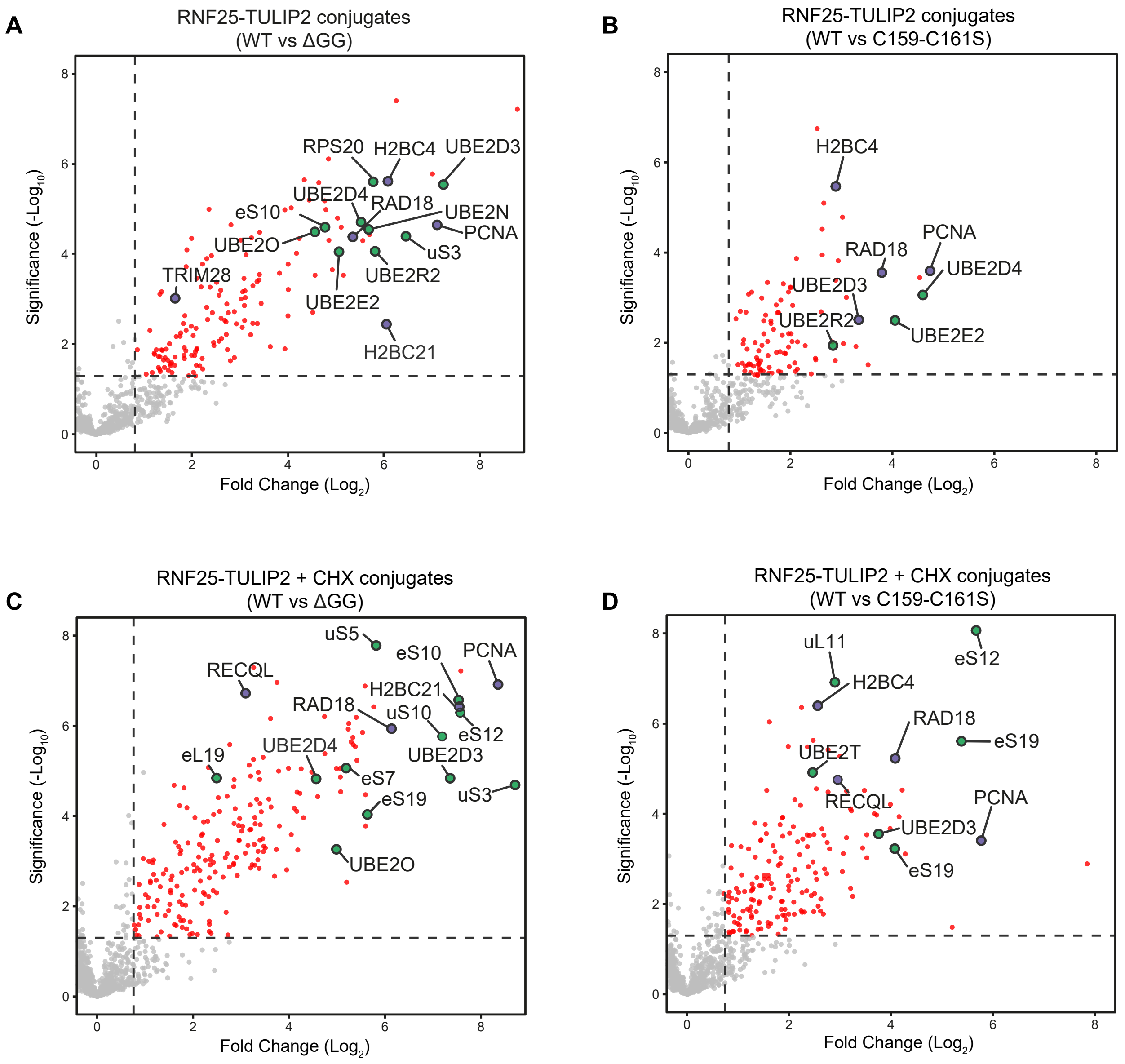


**Supplementary Figure 3.** **(A-D)** Volcano plots depicting statistical differences between RNF25-WT-TULIP2 versus **(A)** RNF25-TULIP2-ΔGG and **(B)** RNF25-C159/161S-TULIP2, and **(C-D)** treated with CHX. Each dot represents a protein, and selected proteins are labeled. Proteins related to similar biological processes are color matched. Dashed lines mark significance threshold for $p > 0.05$ from unpaired two-tailed t-tests and fold change higher than 0.8 (log2).

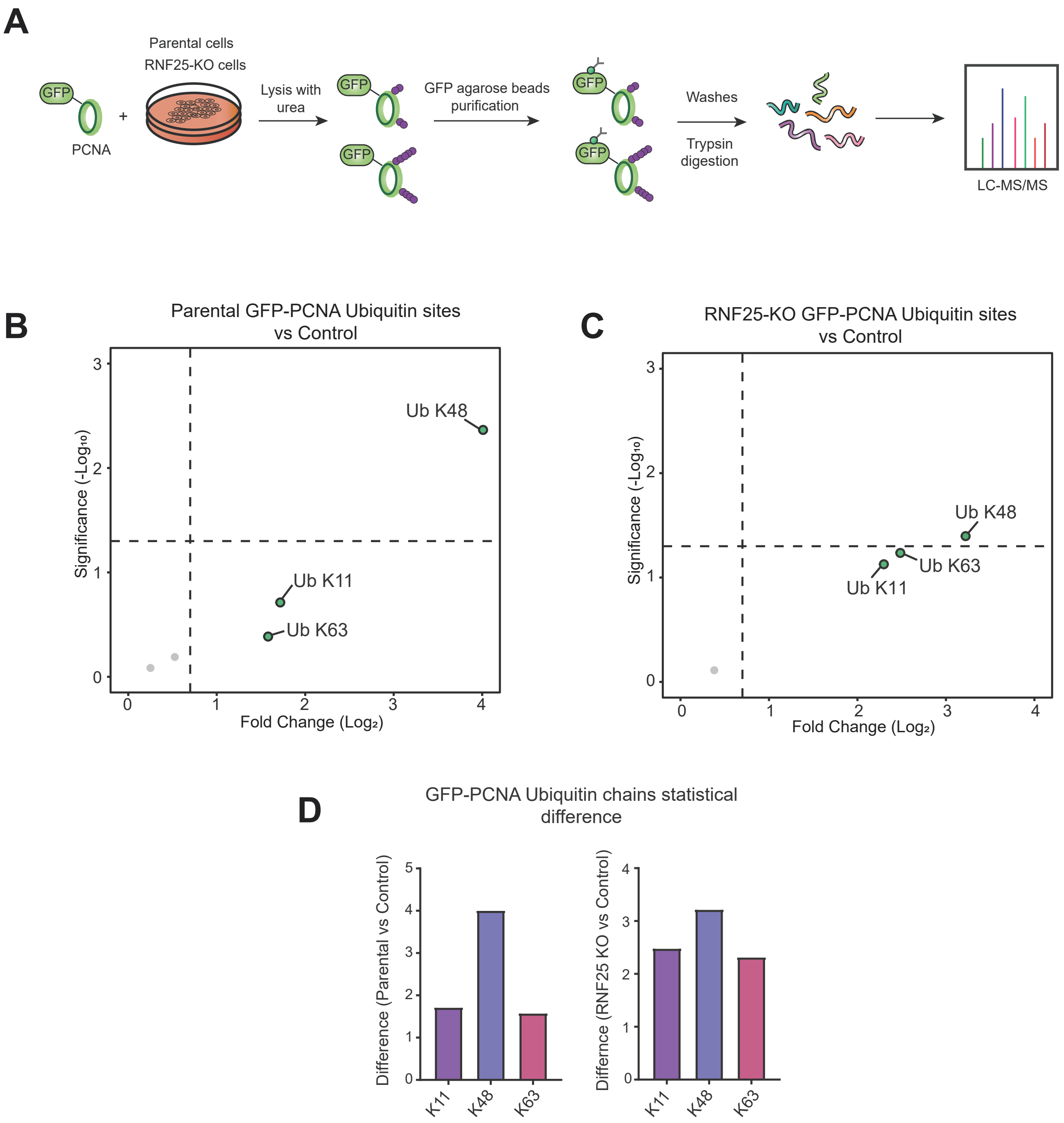


**Supplementary Figure 4. (A)** Illustration of GFP-PCNA immunoprecipitation experiment under denaturing conditions. **(B, C)** Volcano plots showing statistical differences between GlyGlyK sites of **(B)** Parental GFP-PCNA, and **(C)** RNF25-KO GFP-PCNA versus Control. Ubiquitin chains are marked as Ub-lysine (K)-chain number, and ubiquitination lysine sites (K) on specific proteins are labeled. **(D)** Bar plot representing statistical differences regarding ubiquitin chains between Parental GFP-PCNA and Control, and RNF25-KO GFP-PCNA versus Control.

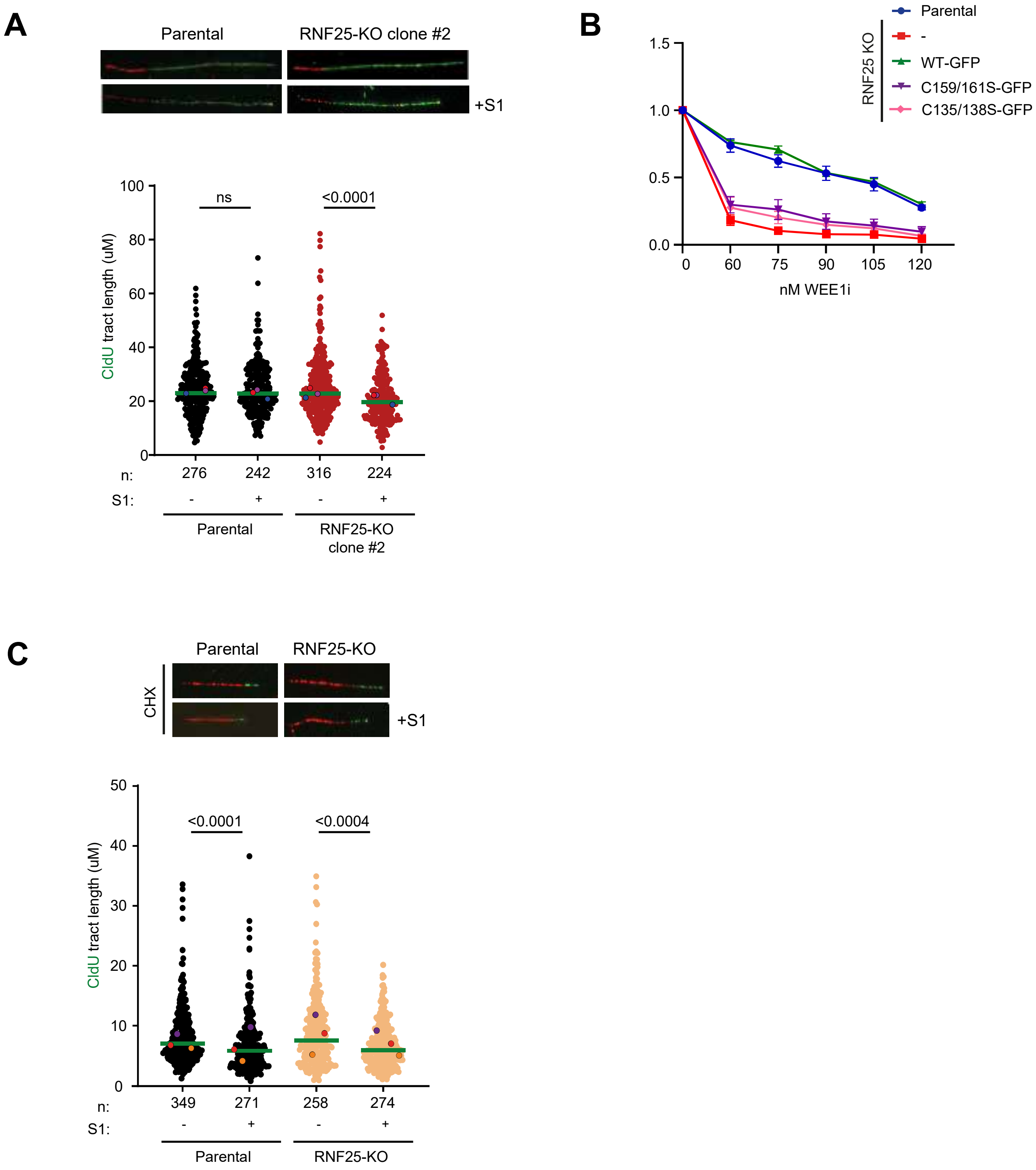


**Supplementary Figure 5. (A)** S1-DNA fiber assay performed with Parental and RNF25-KO clone #2. **(B)** Clonogenic survival assay of Parental and RNF25 KO cells rescued or not with RNF25-WT-GFP, RNF25-C159/161S-GFP or RNF25-C135/138S-GFP under treatment with WEE1 inhibitor for 24 hours and 7-days recovery. Average and standard deviation of three independent experiments with three technical repeats are shown (N = 3). **(C)** S1-DNA fiber assay performed with Parental and RNF25-KO under CHX 3 µg/mL treatment for 1 hour. Below every panel of fluorescence microscopy images, graphs depicting CldU tract lengths of each cell line with and without S1 nuclease treatment are shown. Each dot represents one fiber, and colored dots indicate the median of each independent experiment. The green bar represents the median for all three replicates and n indicates the total fiber count per cell line and treatment. p values were calculated using the two-sided Mann-Whitney test.

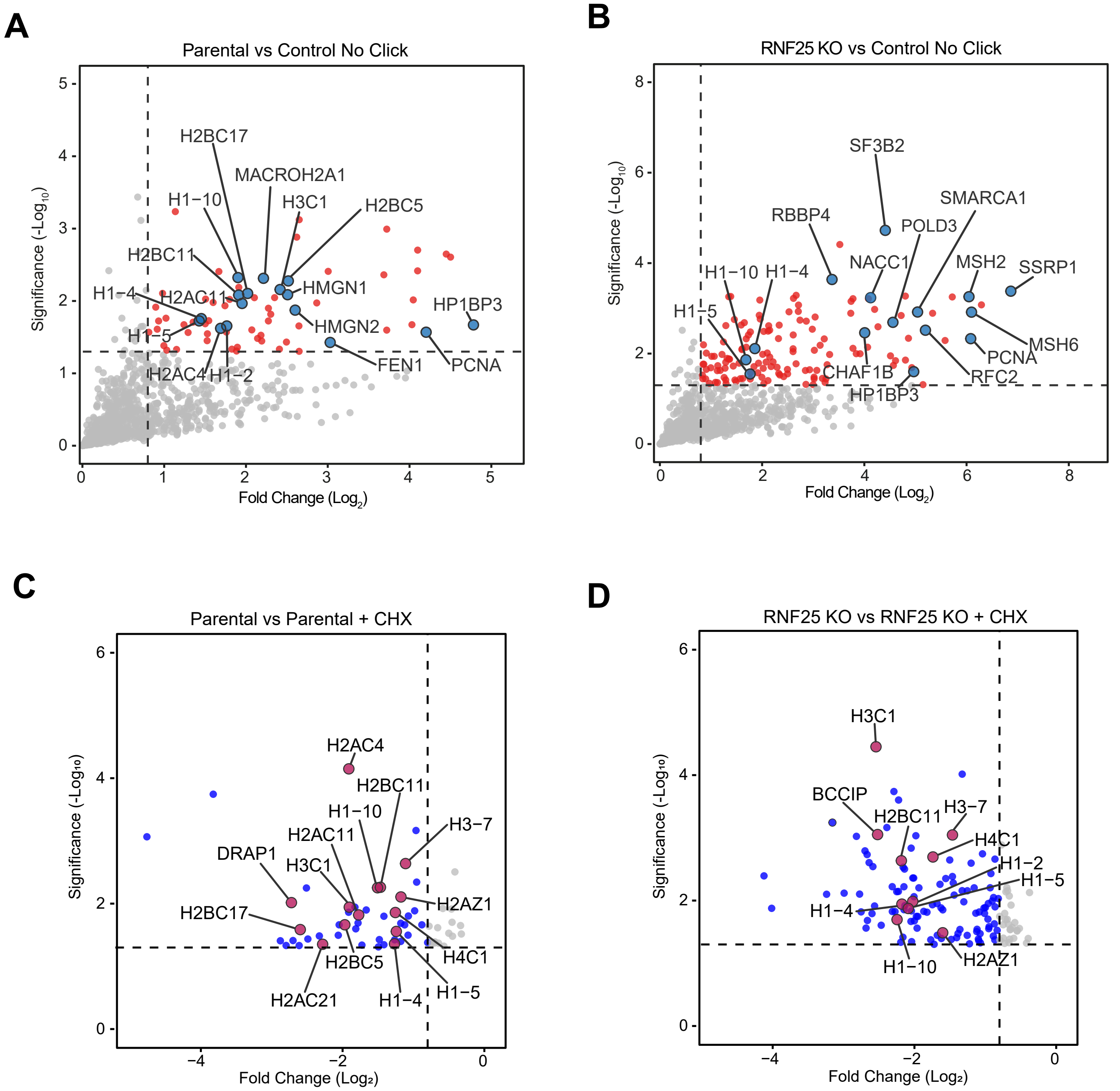


**Supplementary Figure 6. (A, B)** Volcano plots of iPOND experiments depicting **(A)** Parental and **(B)** RNF25-KO replication fork enrichment versus negative control (No Click). Each dot represents a protein. Dotted lines limit significant threshold for a fold-change higher than 0.8 and a -log p higher than 1.3 (p>0.05) for unpaired two-tailed t-tests from biological triplicates. Proteins involved in DNA replication and chromatin organization are highlighted. **(C, D)** Volcano plots of iPOND experiments representing downregulated proteins at the replication fork of **(C)** Parental and **(D)** RNF25-KO cells under CHX treatment, compared to non-treated cells. Each dot represents a protein. Dotted lines limit significant threshold for a fold-change higher than 0.8 and a -log p higher than 1.3 (p>0.05) for unpaired two-tailed t-tests from biological triplicates. Histones proteins are highlighted.

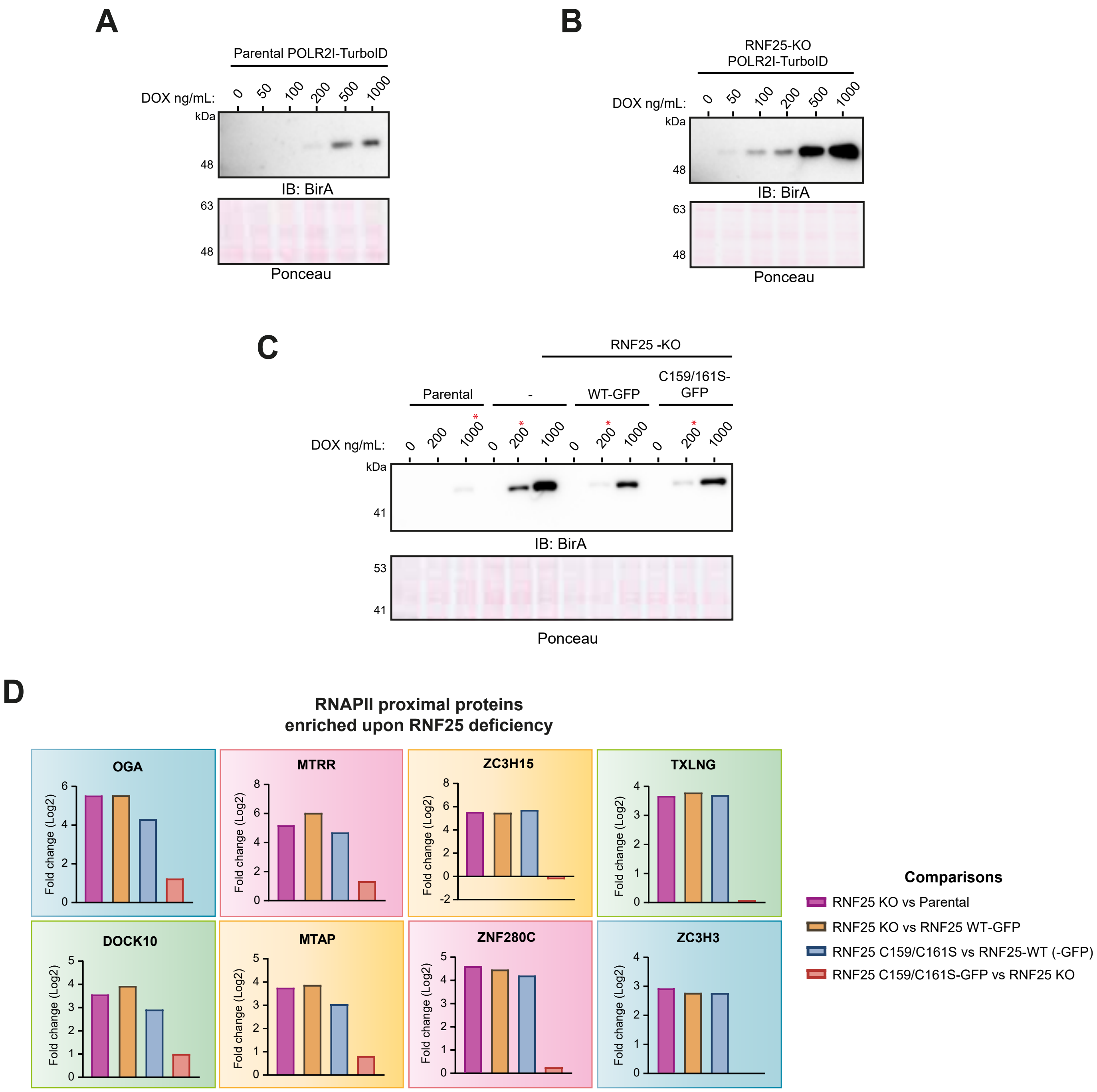


**Supplementary Figure 7. (A-C)** Immunoblotting showing POLR2I-TurboID expression at different doxycycline concentrations in Parental, RNF25-KO, RNF25 WT-GFP and RNF25 C159/161S-GFP cells. POLR2I-TurboID induction concentration used for every cell line is marked with a red asterisk (*). Ponceau staining is shown as loading control. **(D)** Bar graphs depicting the statistical difference between each comparison (fold change) for proteins proximal to RNAPII in a RNF25 mutant background.

**Supplementary Table 1. List of primers**

| **Primer** | **Sequence** | Use |
|---|---|---|
| FW-RNF25-C159S-C161S | CACCCTGTTACCACTACTTCCAC**TCC**CAC**TCC**CTTGCTCGGTACATCCAGCAC | Mutagenesis RING domain RNF25 |
| RV-RNF25-C159S-C161S | GTGCTGGATGTACCGAGCAAG**CCA**GTG**CCA**GTGGAAGTAGTGGTAACAGGGTG | |
| BP-FW-PCNA-GFP | GGGGACAAGTTTGTACAAAAAAGCAGGCTTCATGGTGAGCAAGGGCGA | Gateway BP reaction to introduce GFP-PCNA in pDONR207 |
| BP-RV-PCNA-GFP | GGGGACCACTTTGTACAAGAAAGCTGGGTaCTAAGATCCTTCTTCATCCTCGATCT | |
| FW-sgRNF25-1 | TGGGATTATAGGTGCGAGCC | Target gene knock-out of RNF25 |
| RV-sgRNF25-1 | GCTCAGCCTCAAGGTCAATGA | |
| FW-sgRNF25-2 | AGTAATATTGGTAACGTTAATGC | |
| RV-sgRNF25-2 | CTCCCTTTGCTGCCACACAG | |
| i5 0 | AATGATACGGCGACCACCGAGATCTACACTCTTTCCCTACACGACGCTCTTCCGATCTTTGTGGAAAGGACGAAACACCG | Synthetic Lethality CRISPR screening samples indexes for library preparation |
| i5 1 | AATGATACGGCGACCACCGAGATCTACACTCTTTCCCTACACGACGCTCTTCCGATCTCTTGTGGAAAGGACGAAACACCG | |
| i5 2 | AATGATACGGCGACCACCGAGATCTACACTCTTTCCCTACACGACGCTCTTCCGATC**TGC**TTGTGGAAAGGACGAAACACCG | |
| i5 3 | AATGATACGGCGACCACCGAGATCTACACTCTTTCCCTACACGACGCTCTTCCGATC**TAGC**TTGTGGAAAGGACGAAACACCG | |
| i5 4 | AATGATACGGCGACCACCGAGATCTACACTCTTTCCCTACACGACGCTCTTCCGATC**TCAAC**TTGTGGAAAGGACGAAACACCG | |
| i5 6 | AATGATACGGCGACCACCGAGATCTACACTCTTTCCCTACACGACGCTCTTCCGATC**TTGCACC**TTGTGGAAAGGACGAAACACCG | |
| i5 7 | AATGATACGGCGACCACCGAGATCTACACTCTTTCCCTACACGACGCTCTTCCGATC**TACGCAAC**TTGTGGAAAGGACGAAACACCG | |
| i5 8 | AATGATACGGCGACCACCGAGATCTACACTCTTTCCCTACACGACGCTCTTCCGATC**TGAAGACCC**TTGTGGAAAGGACGAAACACCG | |
| i7 A01 | CAAGCAGAAGACGGCATACGAGAT**CGGTTCAA**GTGACTGGAGTTCAGACGTGTGCTCTTCCGATCTCCAATTCCCACTCCTTTCAAGACCT | |
| i7 B01 | CAAGCAGAAGACGGCATACGAGAT**ATTGGATT**GTGACTGGAGTTCAGACGTGTGCTCTTCCGATCTCCAATTCCCACTCCTTTCAAGACCT | |
| i7 A05 | CAAGCAGAAGACGGCATACGAGAT**ATACTCAA**GTGACTGGAGTTCAGACGTGTGCTCTTCCGATCTCCAATTCCCACTCCTTTCAAGACCT | |

| | | |
|---|---|---|
| i7 A09 | CAAGCAGAAGACGGCATACGAGAT**CGGTGACC**GTGACTGGAGTTCAGACGTGTGCTCTTCCGATCTCCAATTCCCACTCCTTTCAAGACCT | Synthetic Lethality CRISPR screening samples indexes for library preparation |
| i7 B03 | CAAGCAGAAGACGGCATACGAGAT**TATGAGAA**GTGACTGGAGTTCAGACGTGTGCTCTTCCGATCTCCAATTCCCACTCCTTTCAAGACCT | |
| i7 D05 | CAAGCAGAAGACGGCATACGAGAT**GCCACTGG**GTGACTGGAGTTCAGACGTGTGCTCTTCCGATCTCCAATTCCCACTCCTTTCAAGACCT | |
| i7 B10 | CAAGCAGAAGACGGCATACGAGAT**TACAAGTT**GTGACTGGAGTTCAGACGTGTGCTCTTCCGATCTCCAATTCCCACTCCTTTCAAGACCT | |
| i7 C01 | CAAGCAGAAGACGGCATACGAGAT**GCACGACC**GTGACTGGAGTTCAGACGTGTGCTCTTCCGATCTCCAATTCCCACTCCTTTCAAGACCT | |
| i7 H03 | CAAGCAGAAGACGGCATACGAGAT**ATACCTAA**GTGACTGGAGTTCAGACGTGTGCTCTTCCGATCTCCAATTCCCACTCCTTTCAAGACCT | |
| i7 C10 | CAAGCAGAAGACGGCATACGAGAT**ATCAGATT**GTGACTGGAGTTCAGACGTGTGCTCTTCCGATCTCCAATTCCCACTCCTTTCAAGACCT | |

**Supplementary Table 2. List of plasmids**

| Number | Plasmid | Resistance | Origin |
|---|---|---|---|
| pRGP-0001 | HF-TULIP2 | Ampicilin + Chloramphenicol | |
| pRGP-0002 | HF-TULIP2ΔGG | Ampicilin + Chloramphenicol | |
| pRGP-0034 | pLVU-GFP | Ampicilin + Chloramphenicol | Addgene #24177 |
| pRGP-0043 | pX458 | Ampicilin | Addgene #48138 |
| pRGP-0123 | pBLUNT-cut Open | Kanamycin | |
| pRGP-0175 | pDNOR223-RNF25 | Spectinomycin | cDNA ORF library CD0306 |
| pRGP-0180 | pDNOR223-RNF25-C159S-C161S | Spectinomycin | Mutagenesis on pRGP-0175 |
| pRGP-0250 | pLX303 | Ampicilin + Chloramphenicol | Addgene #25897 |
| PRGP-0304 | 10His-ubiquitin-WT-IRES-puro | Ampicilin | |
| pRGP-0329 | sgRNA-RNF25-1 | Ampicilin | |
| pRGP-0330 | sgRNA-RNF25-2 | Ampicilin | |
| pRGP-0331 | HF-RNF25-TULIP2 | Ampicilin | LR pRGP-0175 + pRGP-0001 |
| pRGP-0332 | HF-RNF25-TULIP2ΔGG | Ampicilin | LR pRGP-0175 + pRGP-0002 |
| pRGP-0333 | HF-RNF25-C159S-C161S-TULIP2 | Ampicilin | LR pRGP-0180 + pRGP-0001 |
| pRGP-0354 | pLV-RNF25-WT-GFP | Ampicilin | LR pRGP-0175 + pRGP-0034 |
| pRGP-0358 | pLV-RNF25-C159S-C161S-GFP | Ampicilin | LR pRGP-0180 + pRGP-0034 |
| pRGP-0385 | pCS2-PCNA-GFP | Ampicilin | Gift from Caren Norden (Addgene #105937) |
| pRGP-0386 | pDNR223-POLR2I | Spectinomycin | |
| pRGP-0398 | pDNR223-HIST1H2BN | Spectinomycin | |
| pRGP-0405 | HF-Gateway-GGS-TurboID | Ampicilin + Chloramphenicol | |
| pRGP-0482 | HF-POLR2I-GGS-TURBOID | Ampicilin | LR pRGP-0386 + pRGP-0405 |
| pRGP-0489 | pDNOR207-GFP-PCNA | Gentamycin | pRGP-0385 PCR + BP on pRGP-0047 |
| pRGP-0498 | plx303-GFP-PCNA | Ampicilin | LR pRGP-0489 + pRGP-0250 |
| pRGP-0504 | Gateway-puro | Ampicilin + Chloramphenicol | |
| PRGP-0523 | Gateway-GGS-TurboID-puro | Ampicilin + Chloramphenicol | |
| pRGP-0556 | GGS-TurboID-RNF25 WT | Ampicilin | LR pRGP-0175 + pRGP-0523 |
| pRGP-0557 | GGS-TurboID-RNF25 C159S-C161S | Ampicilin | LR pRGP-0180 + pRGP-0523 |

| | | | |
|---|---|---|---|
| pRGP-0577 | pENTRY100-TurboID start codon | Kanamycin | |
| pRGP-0584 | TurboID only | Ampicilin | LR pRGP-0504 + pRGP-0577 |
| pRGP-0586 | pTSB27-NA85-pLPT-M27-RNH1-EGFP | Ampicilin | Gifted by Emilia Herrera/Andrés Aguilera |

**Supplementary Table 3. Primary antibodies**

| Antibody | Target | Dilution | Company | Catalog number | RRID |
|---|---|---|---|---|---|
| RNF25 | RNF25 | 1:1000 WB | Abcam | ab140514 | AB_2942087 |
| RNF25 | RNF25 | 1:1000 WB | Gifted by Raimundo Freire | | |
| Rad18 (D2B8) | RAD18 | 1:1000 WB | Cell Signaling Technology | 9040 | AB_2756446 |
| PCNA (PC10) | PCNA | 1:5000 WB | Santa Cruz Biotechnology | Sc-56 | AB_628110 |
| Ubiquityl-PCNA (Lys164) (D5C7P) | PCNA K164 | 1:1000 WB | Cell Signaling Technology | 13439 | AB_2798219 |
| Ubiquityl-Histone H2B (Lys120) (D11) | H2B K120 | 1:1000 WB | Cell Signaling Technology | 5546 | AB_10693452 |
| GFP | GFP | 1:1000 WB; 1:500 IF | Novus Biologicals | NB600-308 | AB_10003058 |
| Anti-phospho-Histone H2A.X (Ser139) | yH2AX | 1:500 IF | Millipore | 05-636 | AB_309864 |
| RECQL | RECQL | 1:1000 WB | Proteintech | 14940-1-AP | AB_3085445 |
| BD™ Purified Mouse Anti-BrdU | BrdU | 1:500 IF | Becton Dickinson | 347580 | AB_10015219 |
| Anti-BrdU antibody [BU1/75 (ICR1)] | BrdU | 1:500 IF | Abcam | 6326 | AB_305426 |

**Supplementary Table 4. Secondary antibodies**

| Antibody | Target | Dilution | Company | Catalog number | RRID |
|---|---|---|---|---|---|
| HRP-conjugated Goat-anti-mouse | Mouse IgG (H+L) | 1:10,000 WB | Jackson Immunoresearch | 115-035-146 | AB_2307 392 |
| HRP-conjugated Goat-anti-rabbit | Rabbit IgG (H+L) | 1:10,000 WB | Jackson Immunoresearch | 111-035-003 | AB_2313 567 |
| Goat anti-rat Alexa Fluor 488 | Rat primary antibody IgG | 1:500 IF | Invitrogen | A11006 | AB_2534074 |
| Goat anti-mouse Alexa Fluor 594 | Mouse primary antibody IgG | 1:500 IF | Invitrogen | A11005 | AB_2534073 |
| Goat anti-rabbit Alexa Fluor 488 | Rabbit primary antibody IgG | 1:500 IF | Invitrogen | A11008 | AB_143165 |

### Description of Supplementary Datasets

**Supplementary Dataset 1.** Proteomics data analysis from Parental and RNF25-KO cells expressing or not His-Ubiquitin with and without MG132 treatment.

**Supplementary Dataset 2.** Proteomics data analysis from RNF25-TULIP2 assay treated or not with CHX. Includes RNF25 Wild Type TULIP2 and ∆GG and C159/161S negative controls.

**Supplementary Dataset 3.** Ubiquitin chains detected from GlyGly(K) proteomics data of RNF25-TULIP2 assay treated or not with CHX. Includes RNF25 Wild Type TULIP2 and ∆GG and C159/161S negative controls.

**Supplementary Dataset 4.** Proteomics data analysis from RNF25-GFP coimmunoprecipitation. Includes Parental control without GFP expression, RNF25 Wild Type-GFP and catalytic dead mutant RNF25 C159/161S-GFP.

**Supplementary Dataset 5.** Comparative analysis from proteomic data of RNF25-TurboID assay. Includes Parental control expressing TurboID only, RNF25 Wild Type-TurboID and catalytic dead mutant RNF25 C159/161S-TurboID.

**Supplementary Dataset 6.** Proteomics data analysis from GFP-PCNA denaturing immunoprecipitacion of Parental and RNF25-KO cells expressing GFP-PCNA. Includes Parental control without GFP expression.

**Supplementary Dataset 7.** Ubiquitin chains detected from GlyGly(K) proteomics data of GFP-PCNA denaturing immunoprecipitacion.

**Supplementary Dataset 8.** Proteomics data analysis from iPOND-MS in Parental and RNF25-KO cells treated or not with CHX. Includes Parental and RNF25-KO No Click-It negative controls.

**Supplementary Dataset 9.** Proteomics data analysis from POLR2I-TurboID assay. Includes Parental control expressing TurboID only, Parental and RNF25-KO rescued or not with RNF25 Wild Type-GFP and C159/161S-GFP expressing POLR2I-TurboID.

**Supplementary Dataset 10.** CRISPR screening genes beta scores organized per rank from Parental and RNF25-KO cells.
